# *Enterococcus faecalis* Influences the Transcription and Metabolism of Pathogenic *Escherichia coli* when Grown in Co-Culture

**DOI:** 10.64898/2025.12.08.692790

**Authors:** J Hogins, S. Mekala, J. Resendiz, S. Vedula, A. Nelapolu, Y. Liu, M. Ganjali, A. Wolfe, P.E. Zimmern, L. Freddolino, L. Reitzer

## Abstract

UTI involves bacterial growth in the disparate environments of the bladder and within uroepithelial cells. The bladder is a low-nutrient environment that nonetheless supports rapid growth. Phylogenetic group B2 *Escherichia coli* (Ec) is frequently isolated from urinary tract infection (UTI) patients. Non-B2 strains have also been isolated and are often co-isolated with *Enterococcus faecalis* (Ef). We characterized the interaction between three co-isolated Ec-Ef pairs and other Ec-Ef combinations in a nutrient-rich medium which was intended to emulate the rapid growth condition of the bladder. In this medium, Ef had little effect on Ec growth but resulted in major transcriptome differences. Ef affected the non-B2 and B2 strains differently. For the non-B2 Ec strains, Ef induced transcript for genes whose products degrade ornithine via putrescine to succinate which is subsequently metabolized by the TCA cycle. For a control B2 Ec strain, Ef induced transcripts for growth rate-associated genes of macromolecular synthesis, and for similar metabolic enzymes, except for those that degrade putrescine to succinate. Unexpectedly, Ef induced transcripts for glyoxylate shunt enzymes in both non-B2 and B2 Ec strains. The bacterial disparate and constantly changing environments during UTI suggest the potential for different types of Ec-Ef interactions. Our results provide evidence for nutrient cross-feeding as one type of Ec-Ef interaction.

## Introduction

Urinary tract infection (UTI), one of the most common human bacterial infections, disproportionately affects adult females and costs an estimated $2 billion annually (1–4). The WHO recognizes UTI as a locus for the development of multidrug resistant bacteria that can impact the outcome of other clinically important diseases treated with the same antibiotics (5). This is because antibiotics are the mainstay of therapy with ∼11.3 million prescriptions annually (6, 7). However, first-line antibiotic therapies are not universally effective; a recent randomized trial reported high failure rates in symptom resolution at 28 days (27-39%) and microbial resolution at 14 days (18-27%)(8). This is consistent with the clinical observation that up to 50% of adult females experience UTI recurrence within 6-12 months (9) and up to 15% of females over 60 years of age develop frequent UTIs (10, 11); of females who experience recurrence, a subgroup (∼2-5%) experience frequent recurrences (1). Most females who meet the diagnostic criteria for recurrent UTI (rUTI) ― defined as culture documented UTI ≥2 per 6 months or ≥3 per year ― undergo evaluation without identifying a clear etiology, such as medical or surgical causes (for example, severe immunosuppression or an intravesical foreign body, respectively).

In people diagnosed with UTI, *Escherichia coli* is most often detected (12, 13), with phylogenetic group B2 strains constituting a majority of the detected *E. coli* (12, 14, 15). Group B2 strains also are associated with a variety of *E. coli*-caused diseases (16–21), are more often associated with colorectal cancer (22), and possess more characteristics associated with multiple antibiotic resistance and extended spectrum β-lactamase production (23). When grown in rich medium, group B2 strains express more transcripts for ribosomal proteins, purine synthesis enzymes, ribonucleotide reductase, DNA replication enzymes, DNA gyrase, polyamine synthesis, and RNA polymerase subunits which, in aggregate, suggest faster growth (24). This supposition is supported by *in vivo* growth rate measurements that suggest *E. coli* in the bladder can exhibit a generation time of only 22 minutes (25). Such rapid growth is thought to be critical as only one in 10^5^-10^6^ cells survive passage through urothelial cells (26), which implies maintenance of a high bacterial cell density and a theoretical study found that a doubling time of about 30 minutes can maintain a high density in the bladder lumen (27).

*E. coli* can be co-isolated with other species with pathogenic potential, for example *Enterococcus faecalis* (28). We recently isolated 36 clinical *E. coli* isolates from patients diagnosed with UTI: *E. faecalis* co-isolated with eight non-B2 group strains but with none of 28 phylogroup B2 strains [(28) and unpublished results). The probability of this co-occurrence is 0.00017, suggesting that *E. coli-E. faecalis* co-occurrence may be related to the *E. coli* phylogroup. While it is too early to state with certainty that this phylogenetic association is a general rule, the interaction of the *E. coli-E. faecalis* pairs can be studied. We therefore co-cultured these *E. coli* and *E. faecalis* strains in a rich medium to simulate the rapid growth observed in the bladder. We chose this rich medium instead of a urine-like environment because the rapid growth rate *in vivo* is not easily replicated *in vitro* due to the difficulty in maintaining a low-nutrient chemostat-like environment. In this rich medium, we found that Ef had little effect on Ec growth but resulted in major transcriptome differences, the Ec-Ef interactions were not strain-specific, Ef affected the B2 strain differently, and Ef affected Ec metabolic gene transcription. The results also suggest the basis for interspecies interactions that might be associated with a variety of diseases whenever one phase of infection involves rapid bacterial growth.

## Results

### The effect of co-culturing on growth

Strains (described in Fig 1A) were grown for two hours in glucose-tryptone (GT) medium inoculated from colonies on species-specific selective media. After two hours, the cultures were diluted at least 100-fold into fresh medium while ensuring equal inoculation densities for each strain. Growth was measured at an optical density of 600 nm every 30 minutes. Ec monocultures grew better than Ef monocultures (Fig 1B). The mixed culture optical densities closely paralleled those of the Ec monocultures, which means that the change in optical density for mixed cultures is more of a reflection of the effect of Ef on Ec than the reverse. In general, Ec is compatible with Ef which appeared to have little effect on Ec growth. Considering the high nutrient content of GT medium, this result is not unexpected because Ec and Ef are not competing for limiting nutrients.

**Fig 1:**
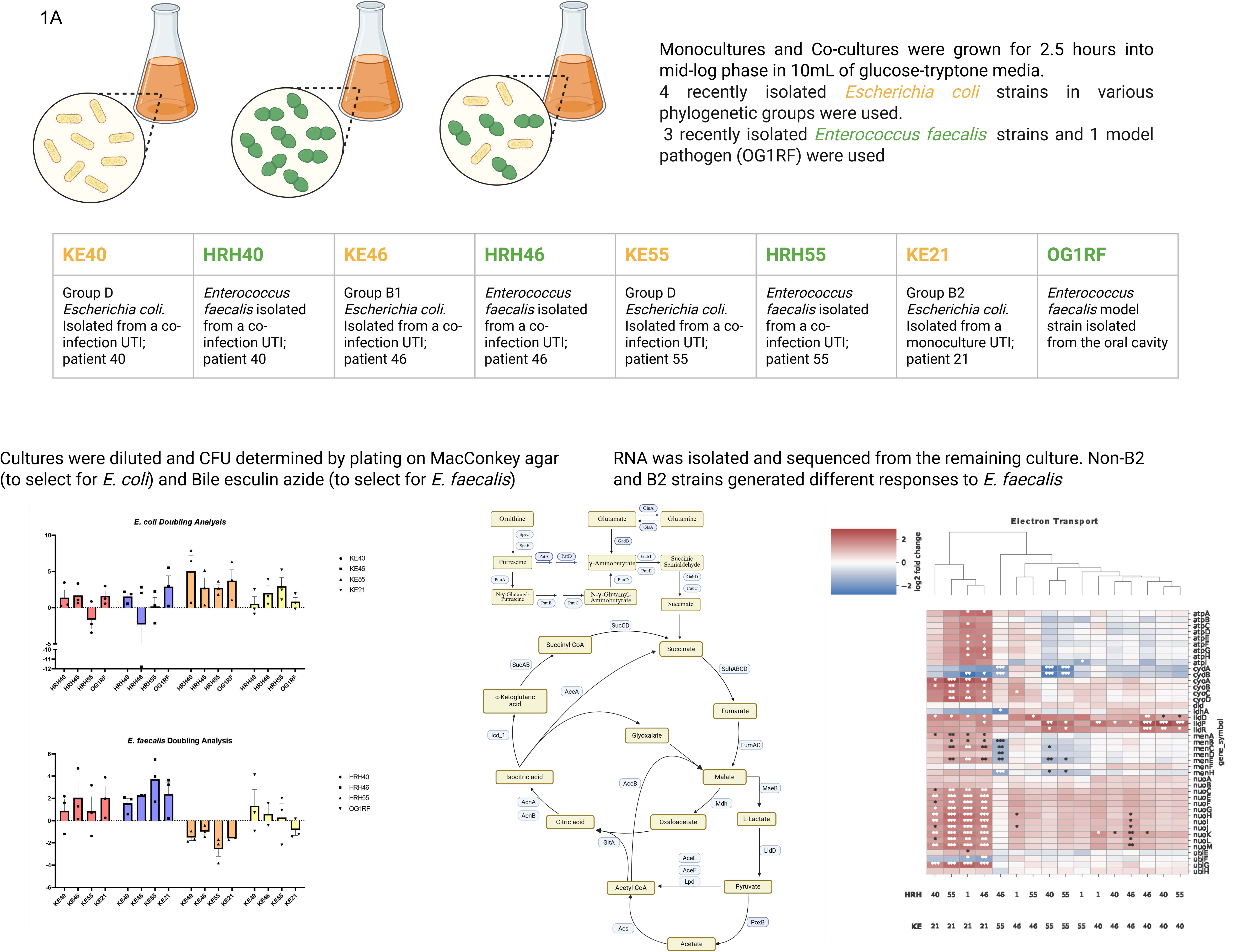
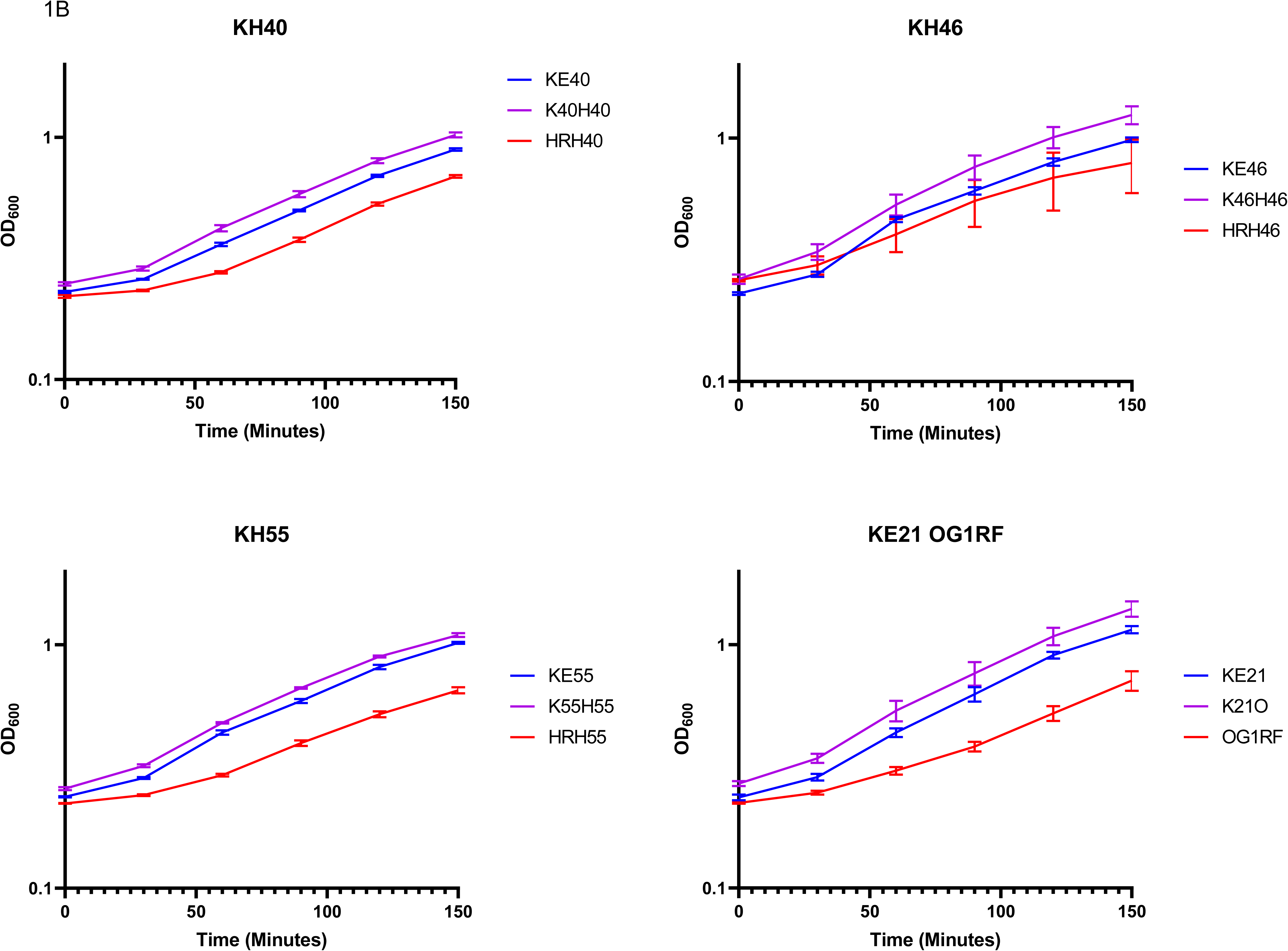
Growth. A) Experimental schematic. B) Growth measures of paired strains. The mono-cultures (Ef in red) (Ec in blue) and co-cultures (purple) are shown. The points represent the average with the standard error mean shown as bars.

### The effect of co-culturing on Ec and Ef transcriptomes

After strains were mixed, cultures were harvested by centrifugation during exponential growth (OD600 of 0.3 to 0.5) for mRNA extraction. We observed profound changes to the transcriptomes of co-cultured strains, which are distinct in UMAP plots from the corresponding monocultures, with typically several hundred transcripts differentially expressed between those conditions. (Fig 2A-C).

**Fig 2:**
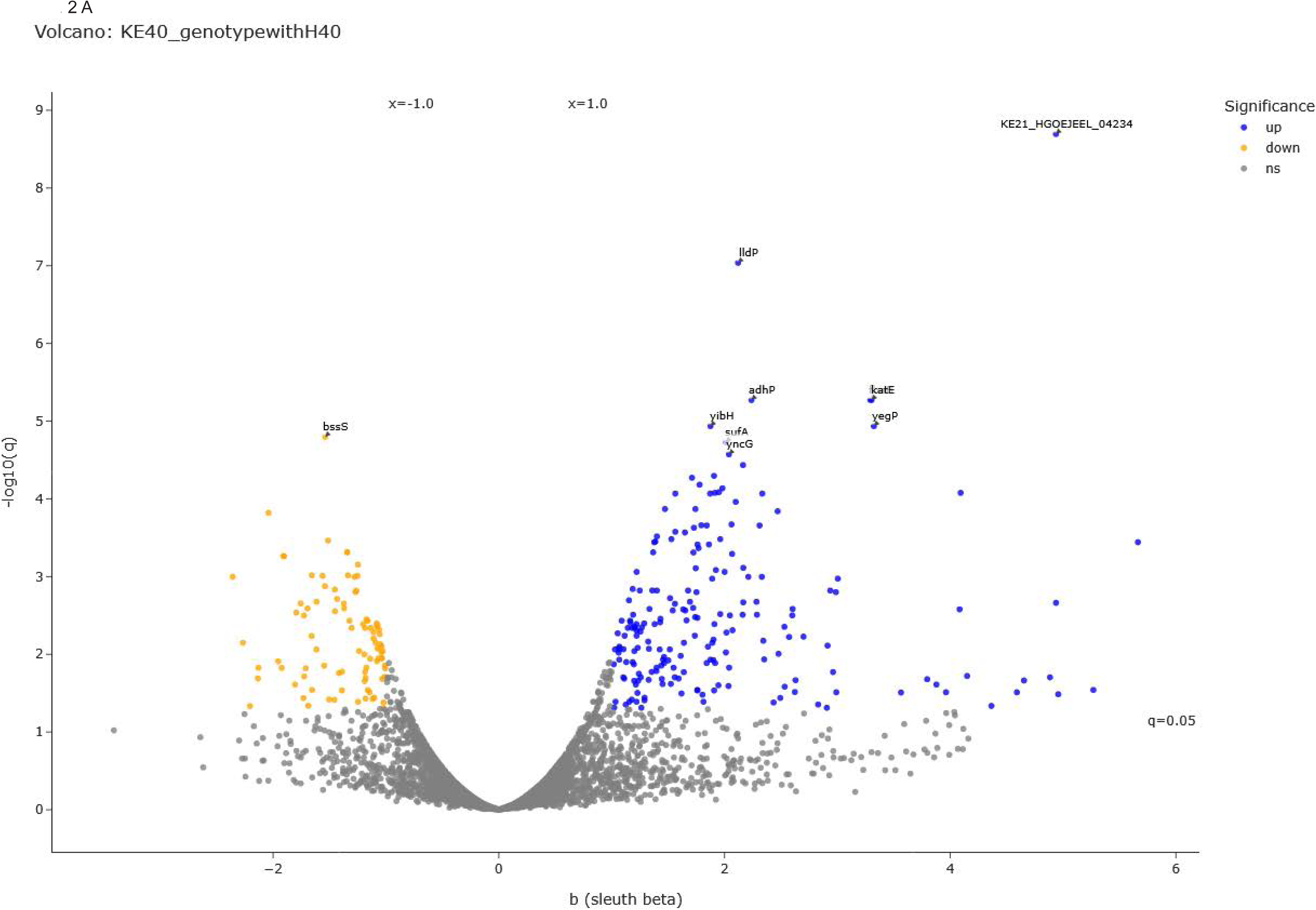
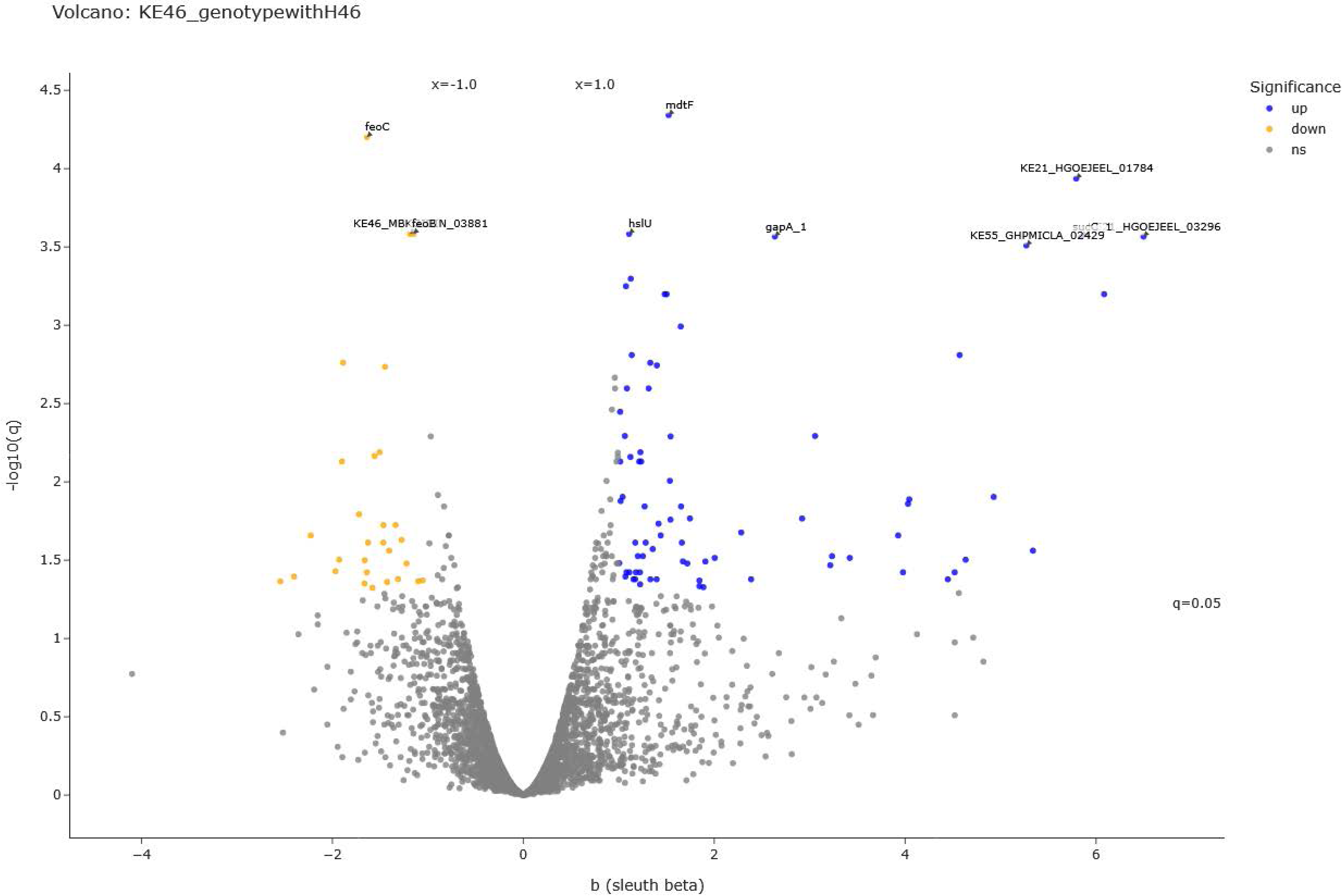
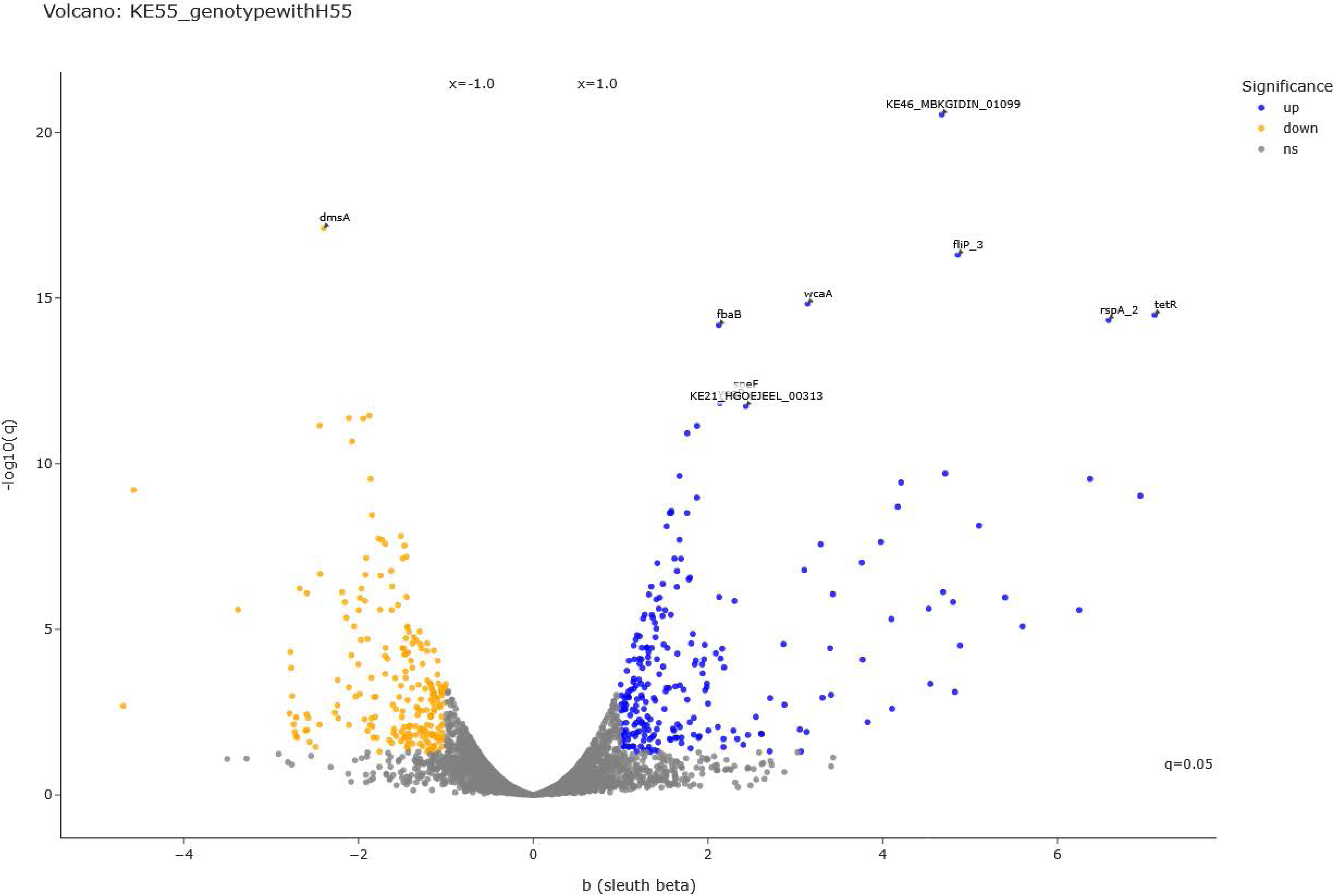
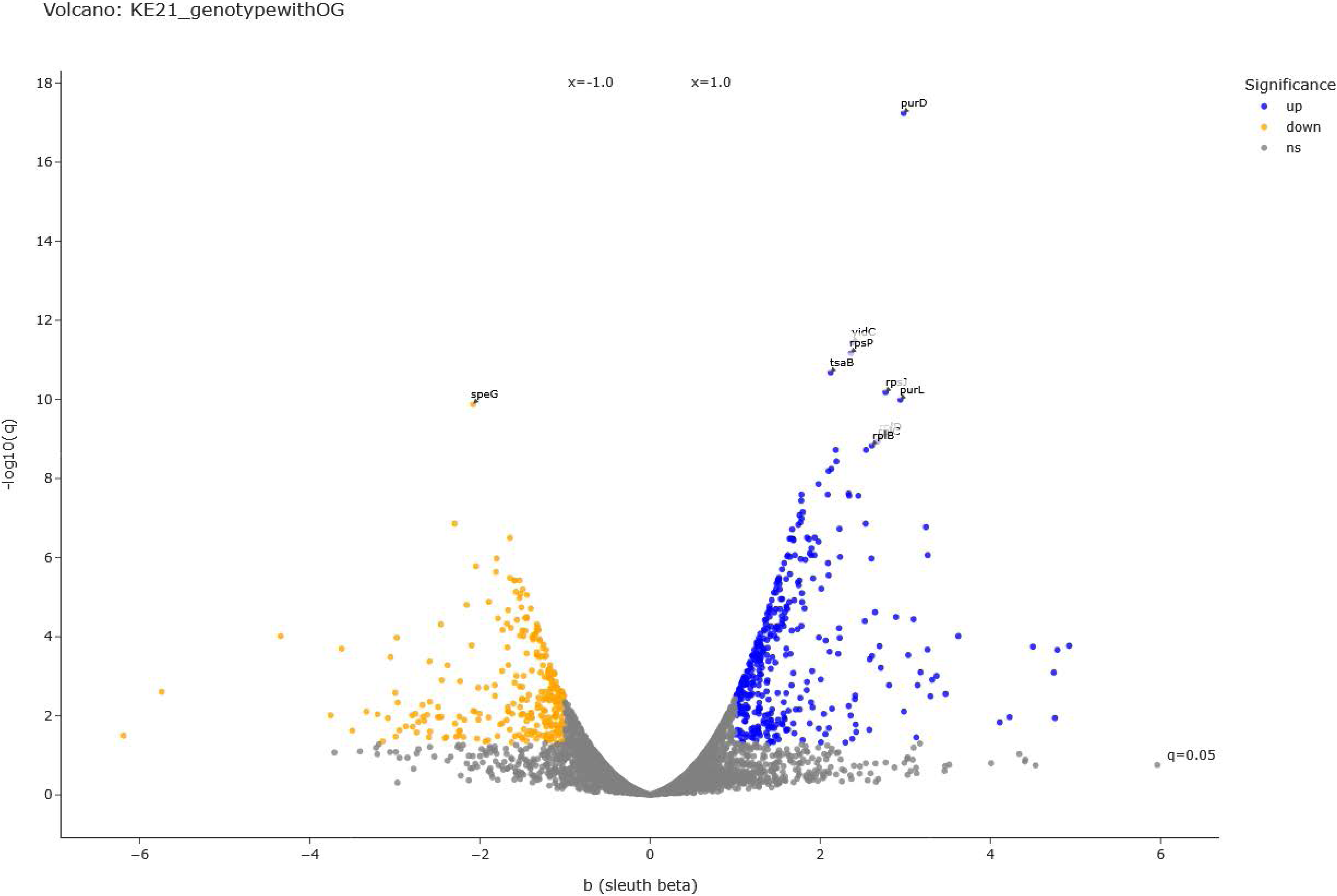
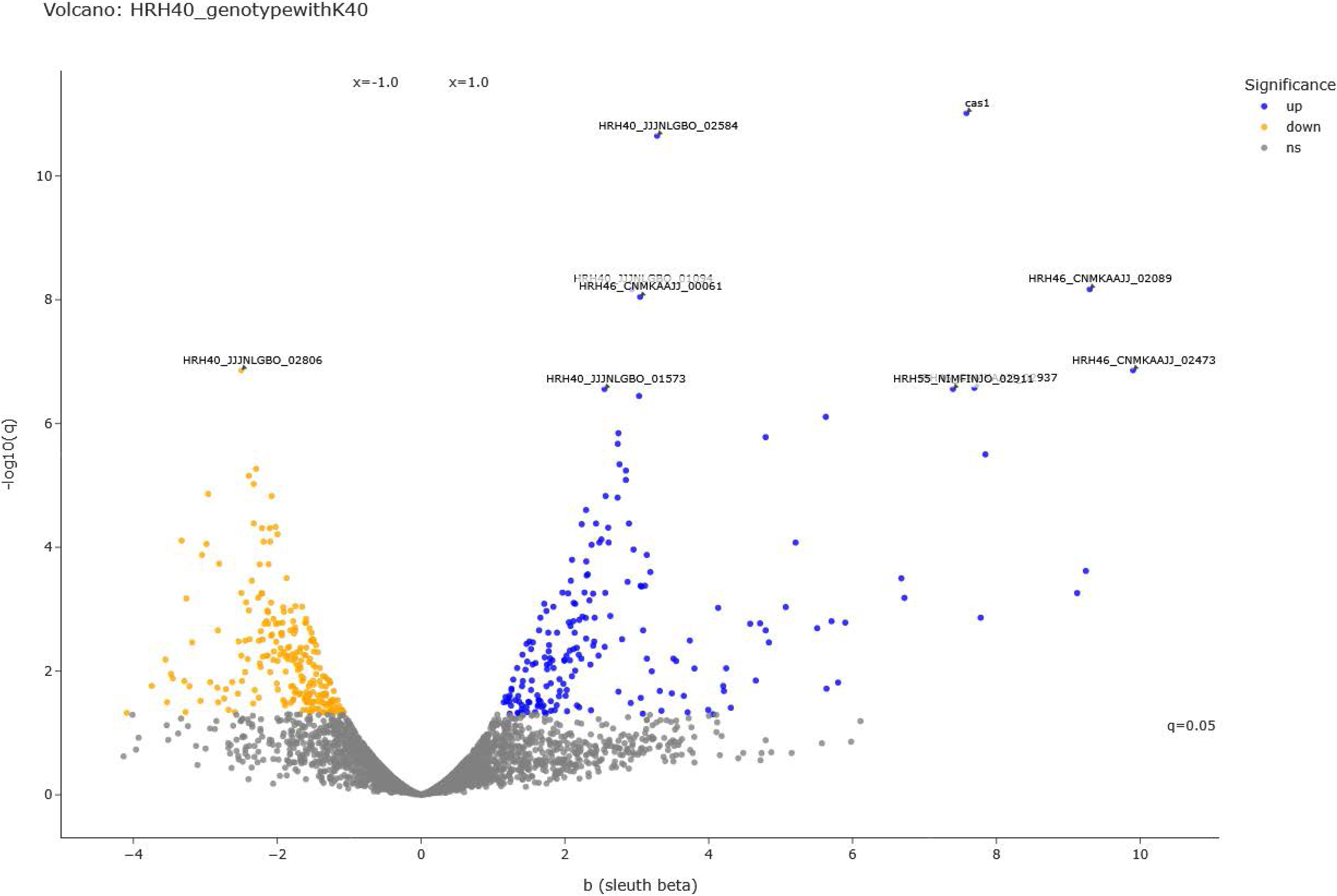
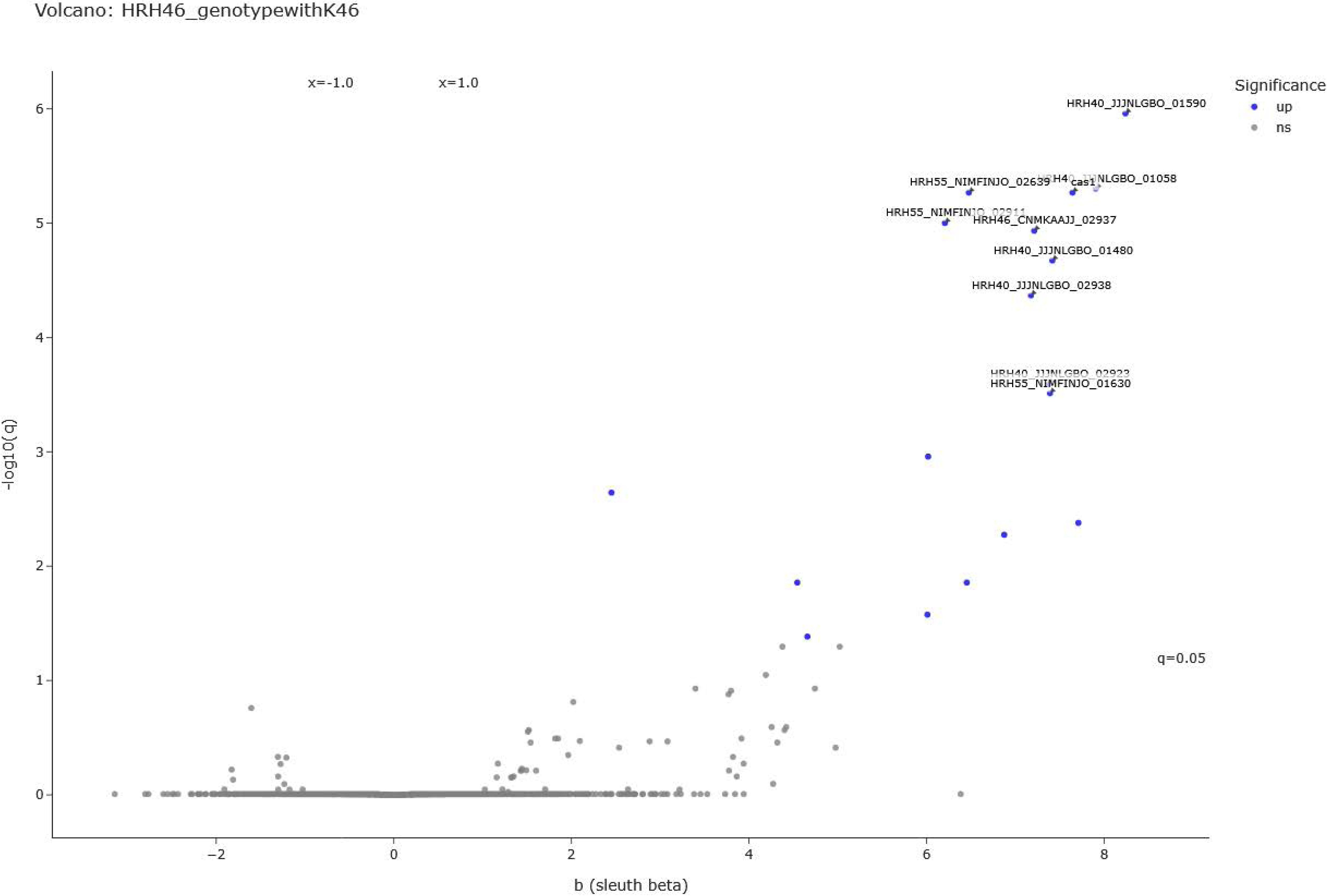
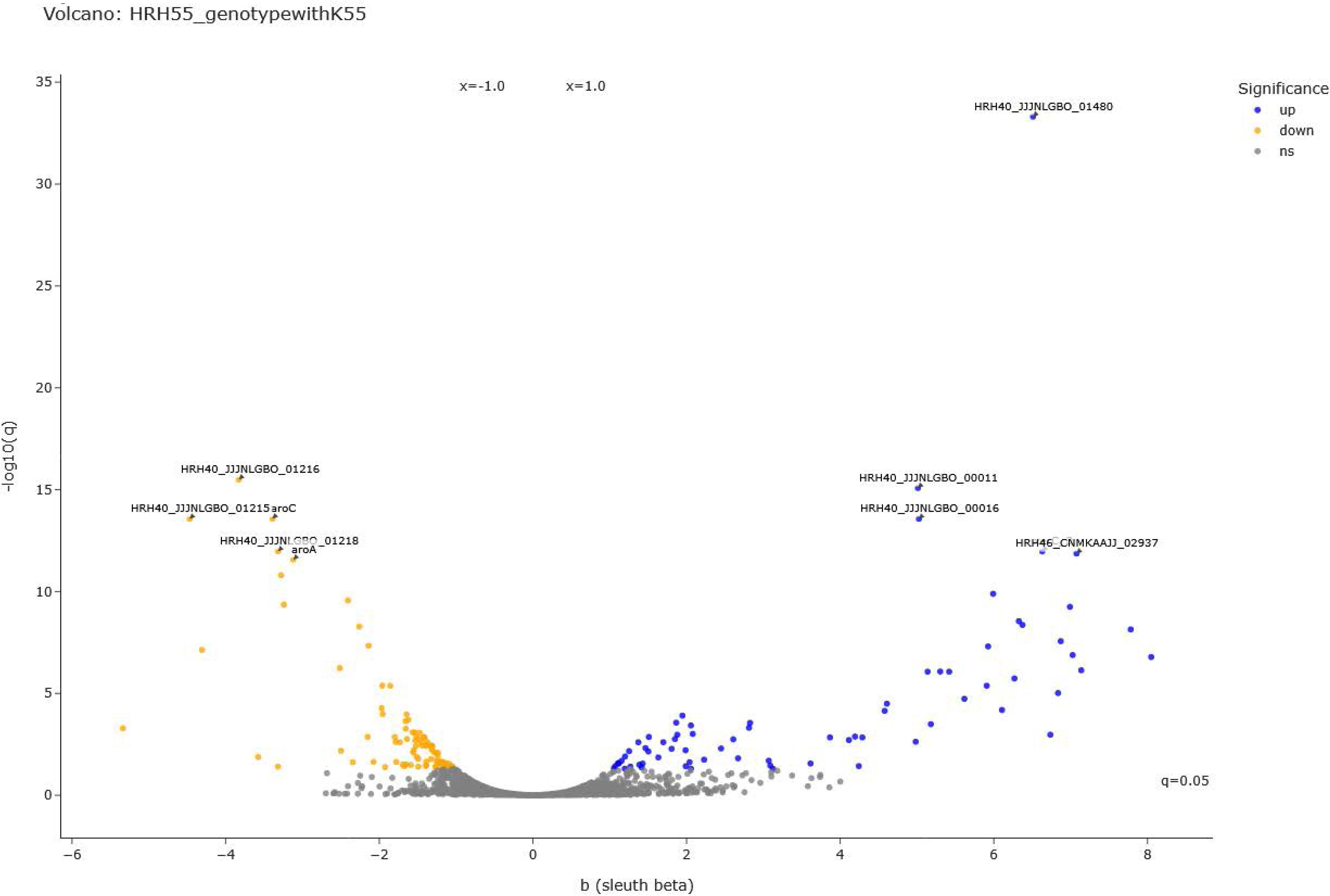
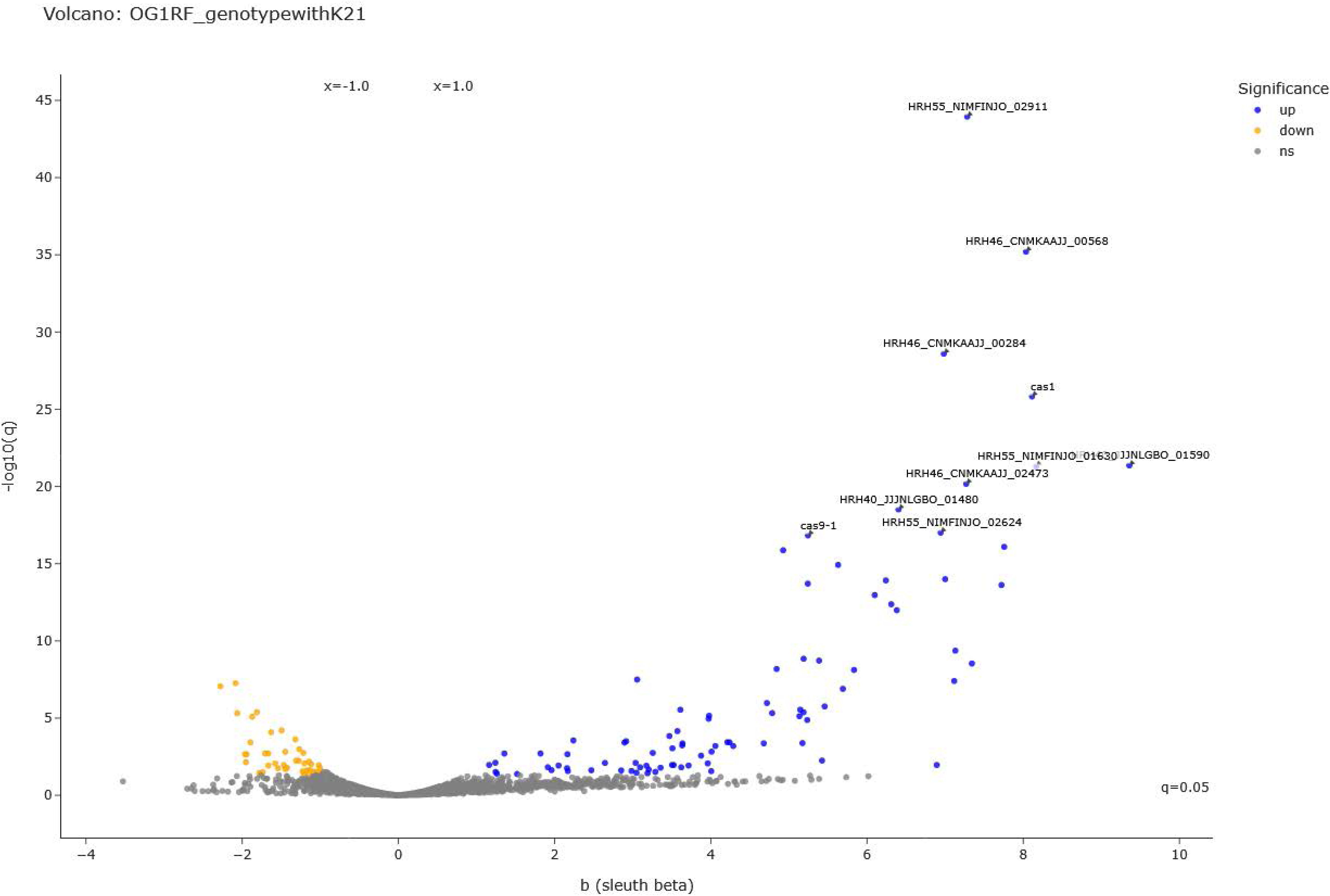
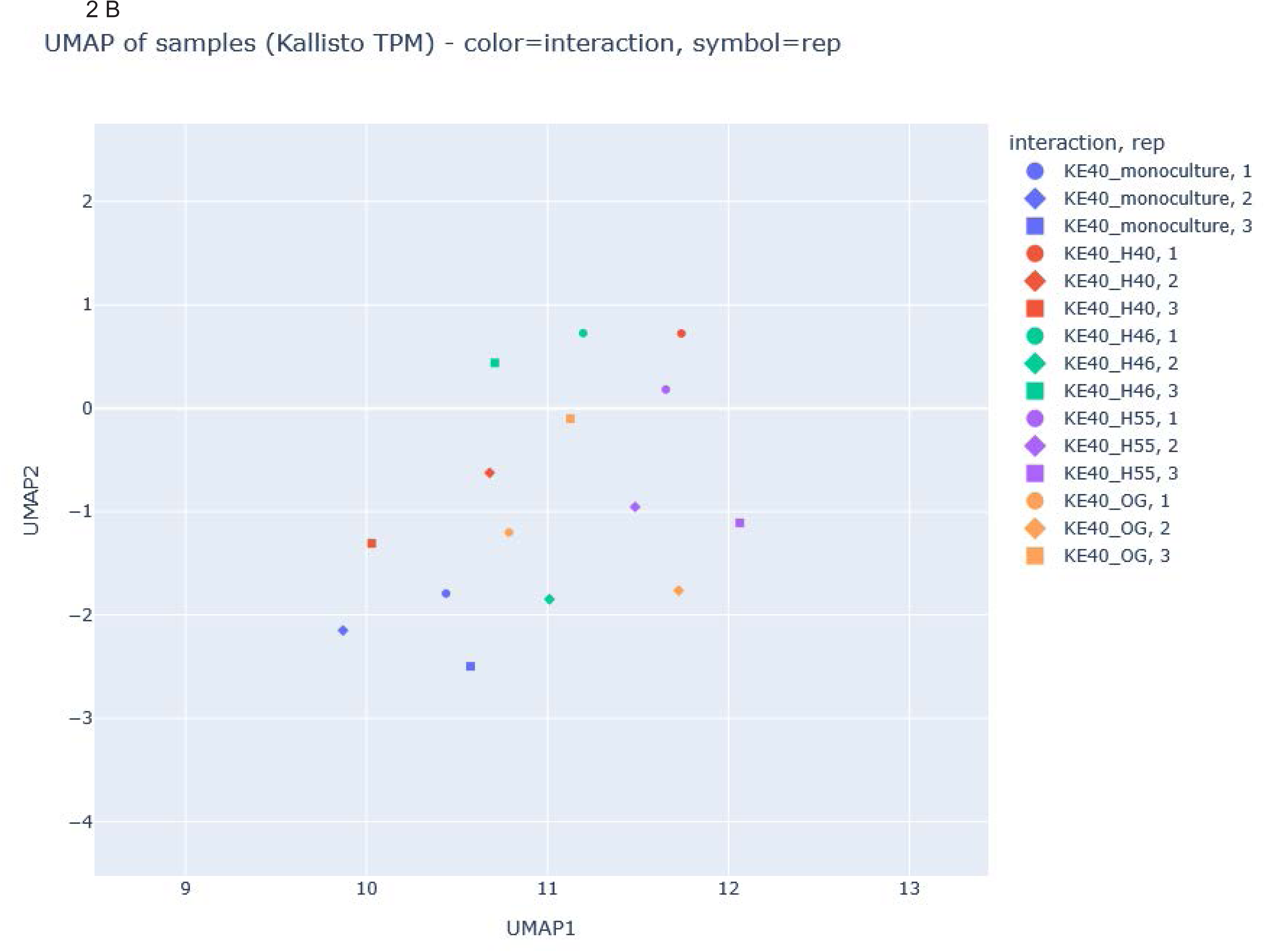
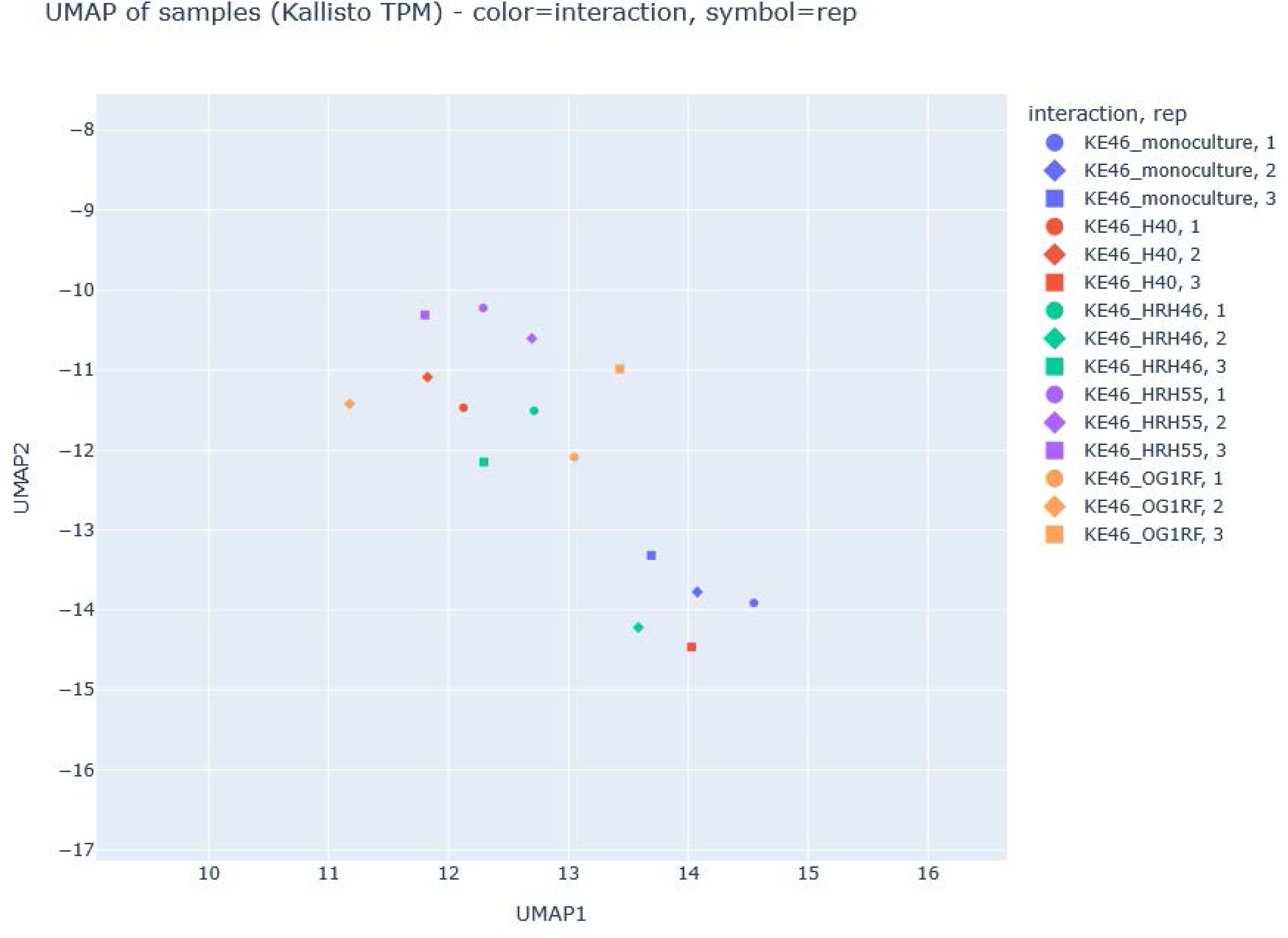
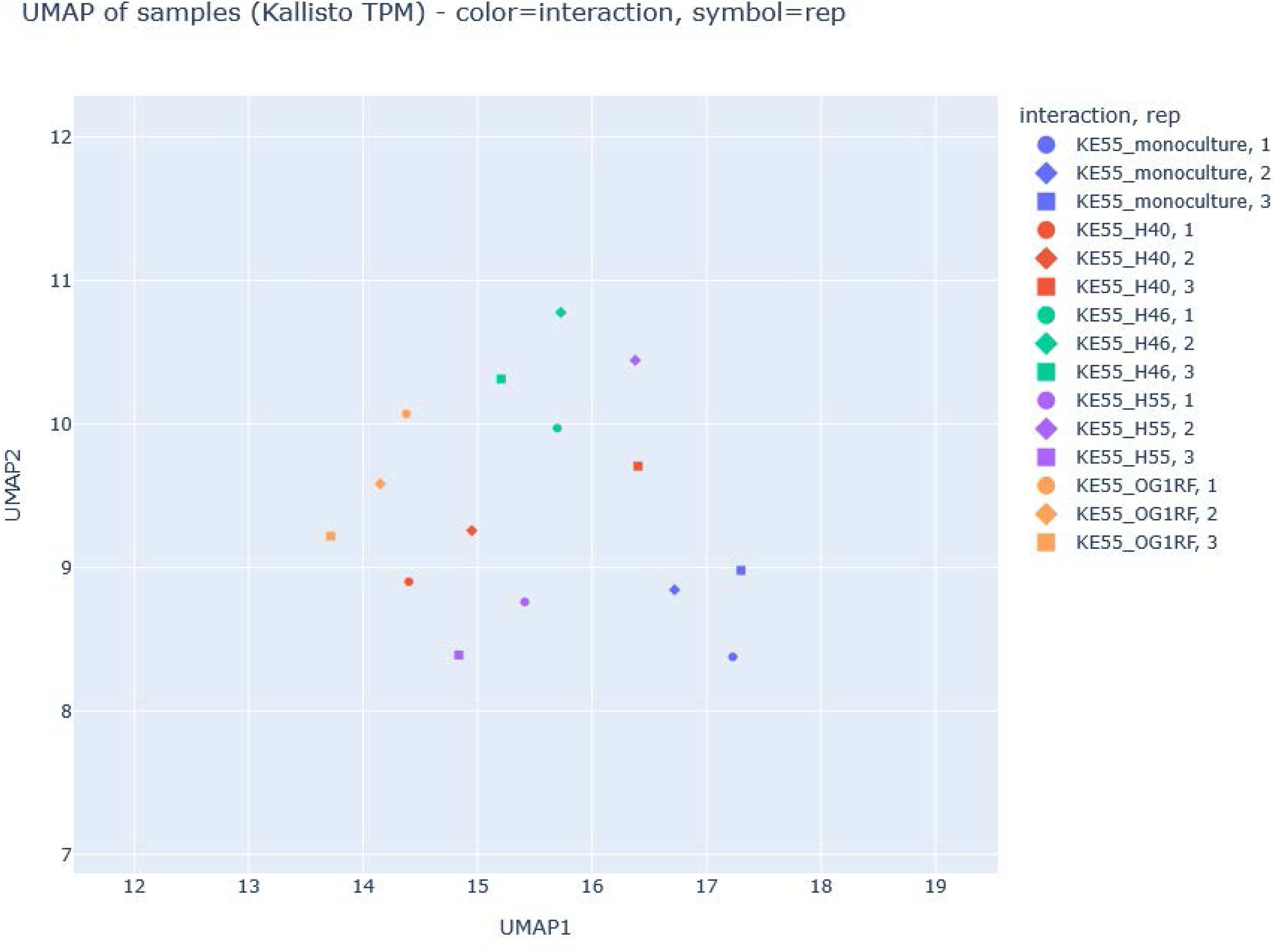
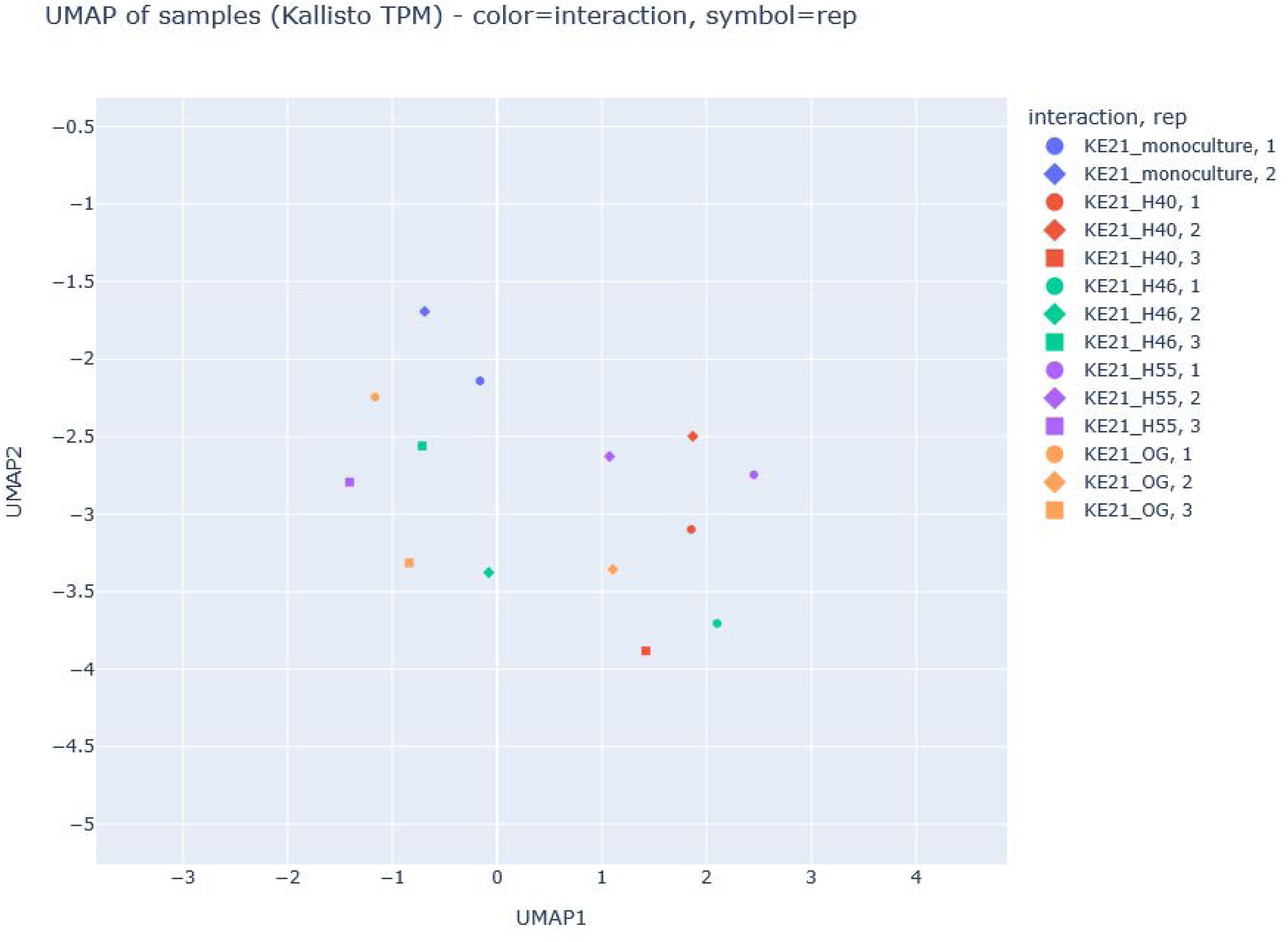
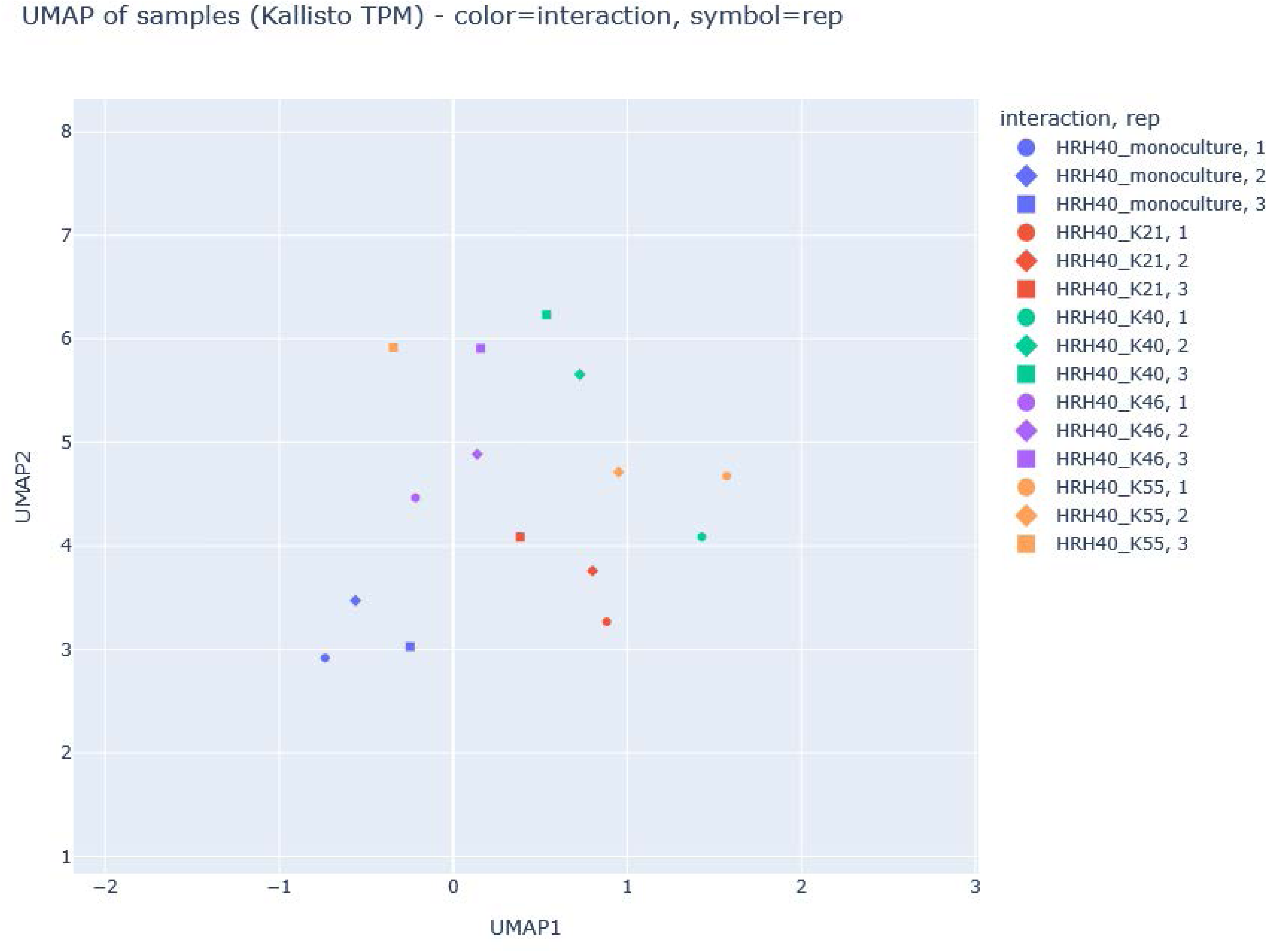
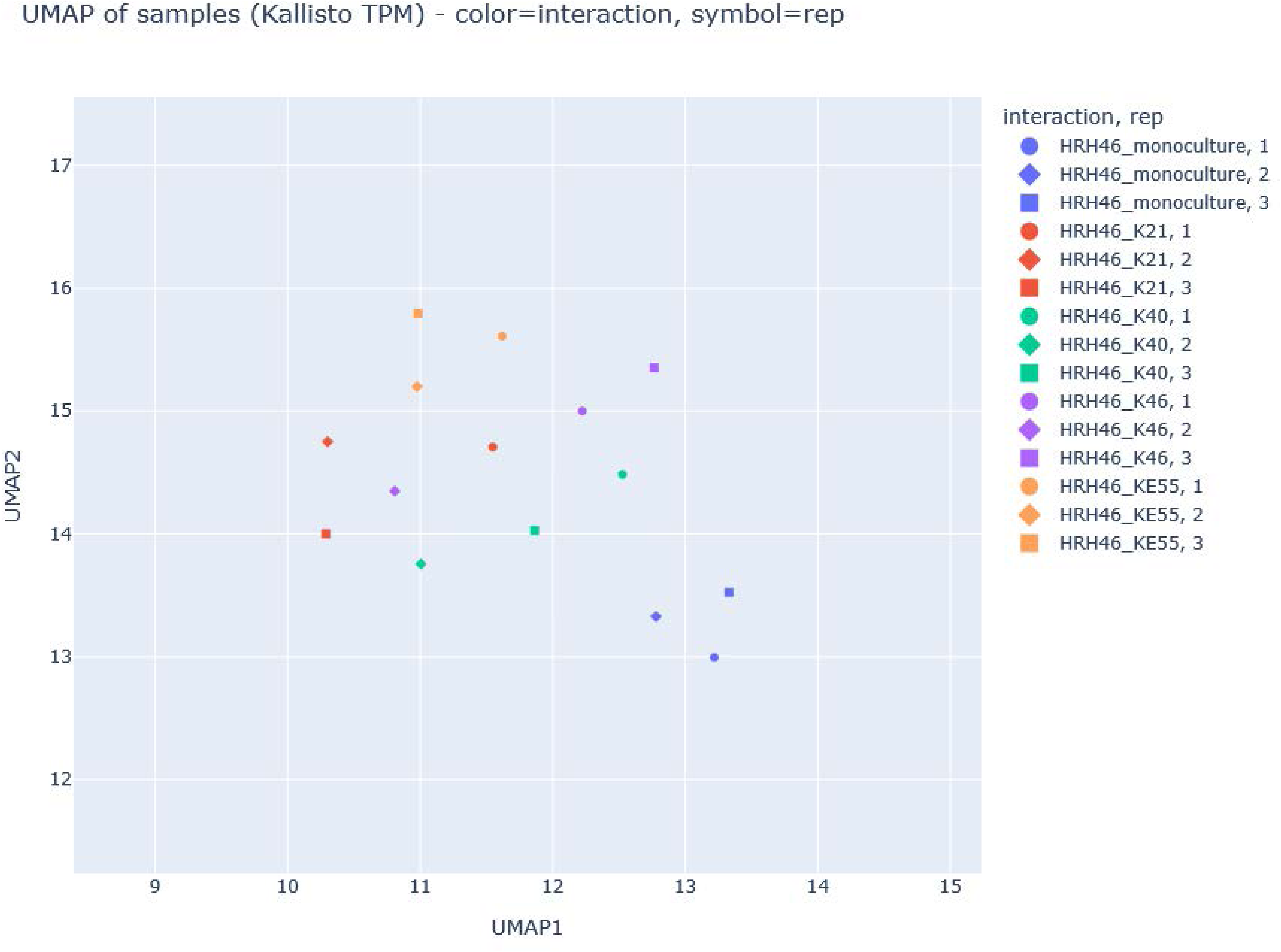
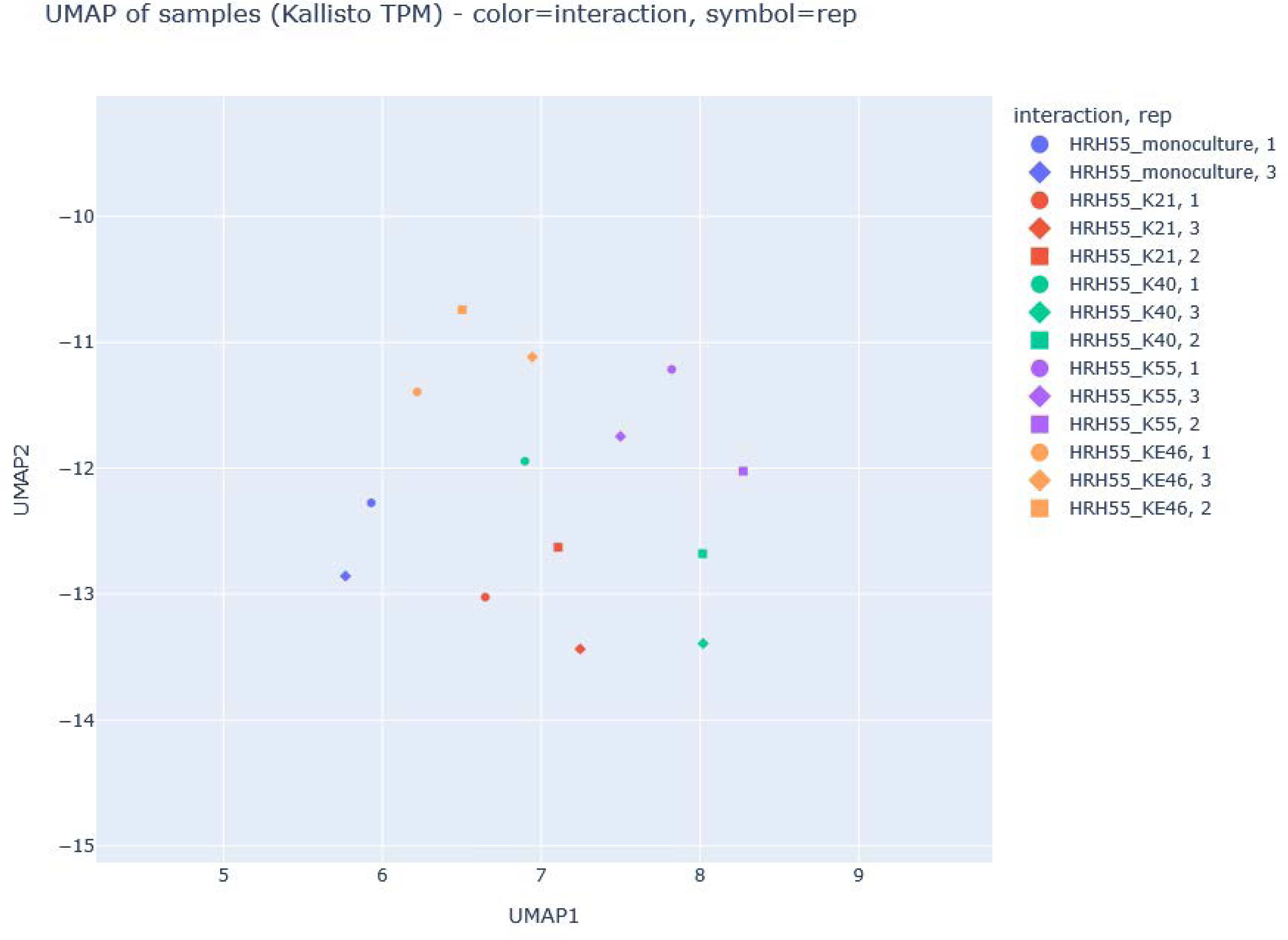
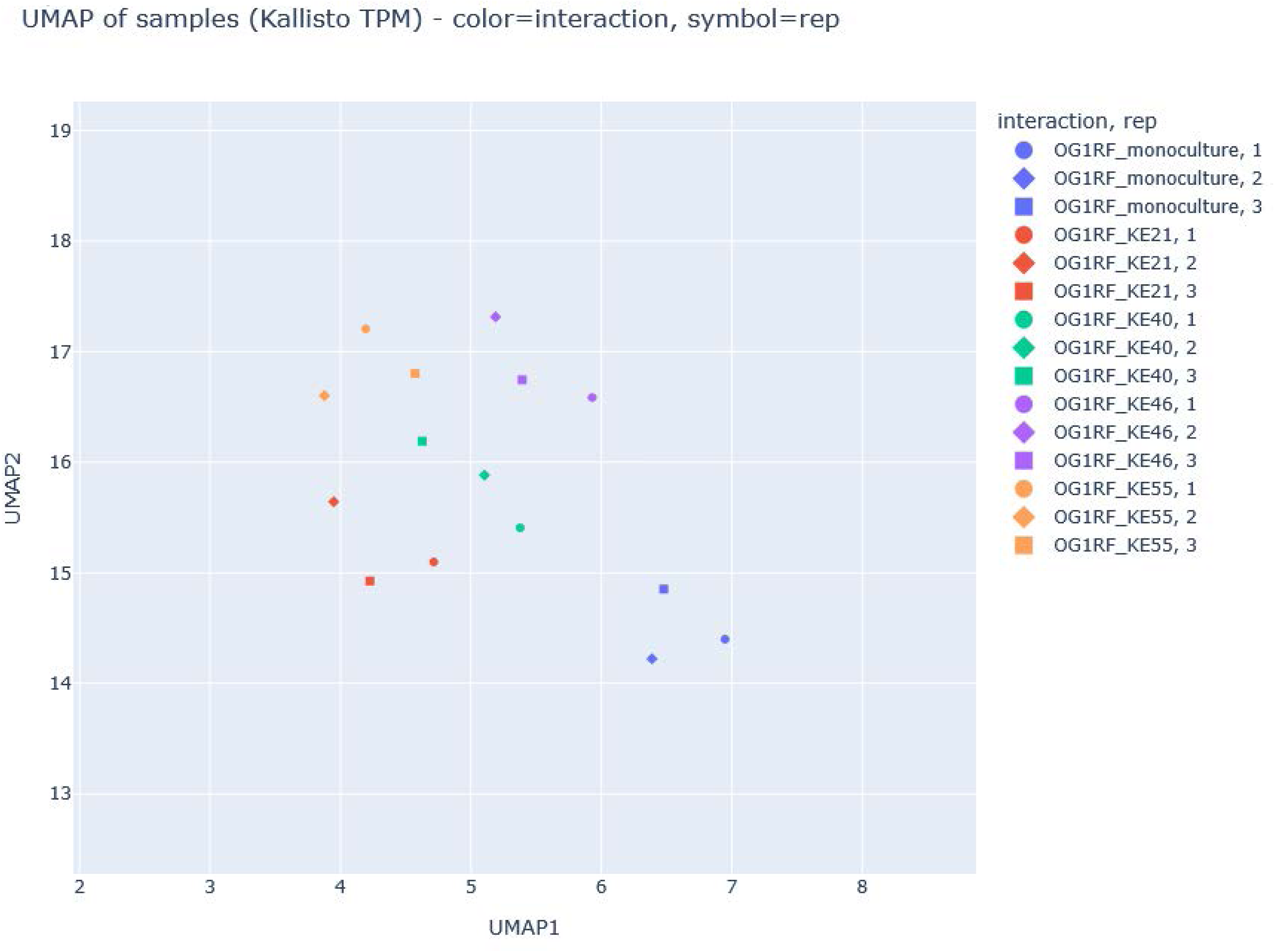
General Transcriptomic Changes. Summary of the transcriptomic data. A) Volcano plots of representative pairs. The Ec strains show large significant transcriptomic changes, while only HRH40 show robust significant changes in the Ef strains. B) UMAP analysis of each replicate for each pairing and mono-culture. The monocultures appear to cluster together while the co-cultures form clusters away from the monoculture cluster.

Gene set enrichment analysis across the co-cultured pairs showed recurring patterns in the *E. coli* transcriptomes during interaction with *E. facaelis*, with particularly prominent and frequent changes in the expression of genes involved in respiration, central carbon metabolism, polyamine biosynthesis, and stress responses (Fig 3). As a particularly prominent example, HRH40 showed significant differential expression of 148, 443, and 244 transcripts (q < 0.05) for interaction with KE40, KE46, and KE55, respectively, which are the largest number of differences, except for the KE55-HRH46 interaction. The other interactions had fewer unique differences. The four Ef strains had 43, 18, and 149 common transcript differences for KE40, KE46, and KE55, respectively, and 348 common transcripts differences for interactions with the B2 strain KE21. In other words, Ef had a greater effect on the B2 strain than on the non-B2 strains, which is consistent with cooperation between Ef and the non-B2 strains.

**Fig 3:**
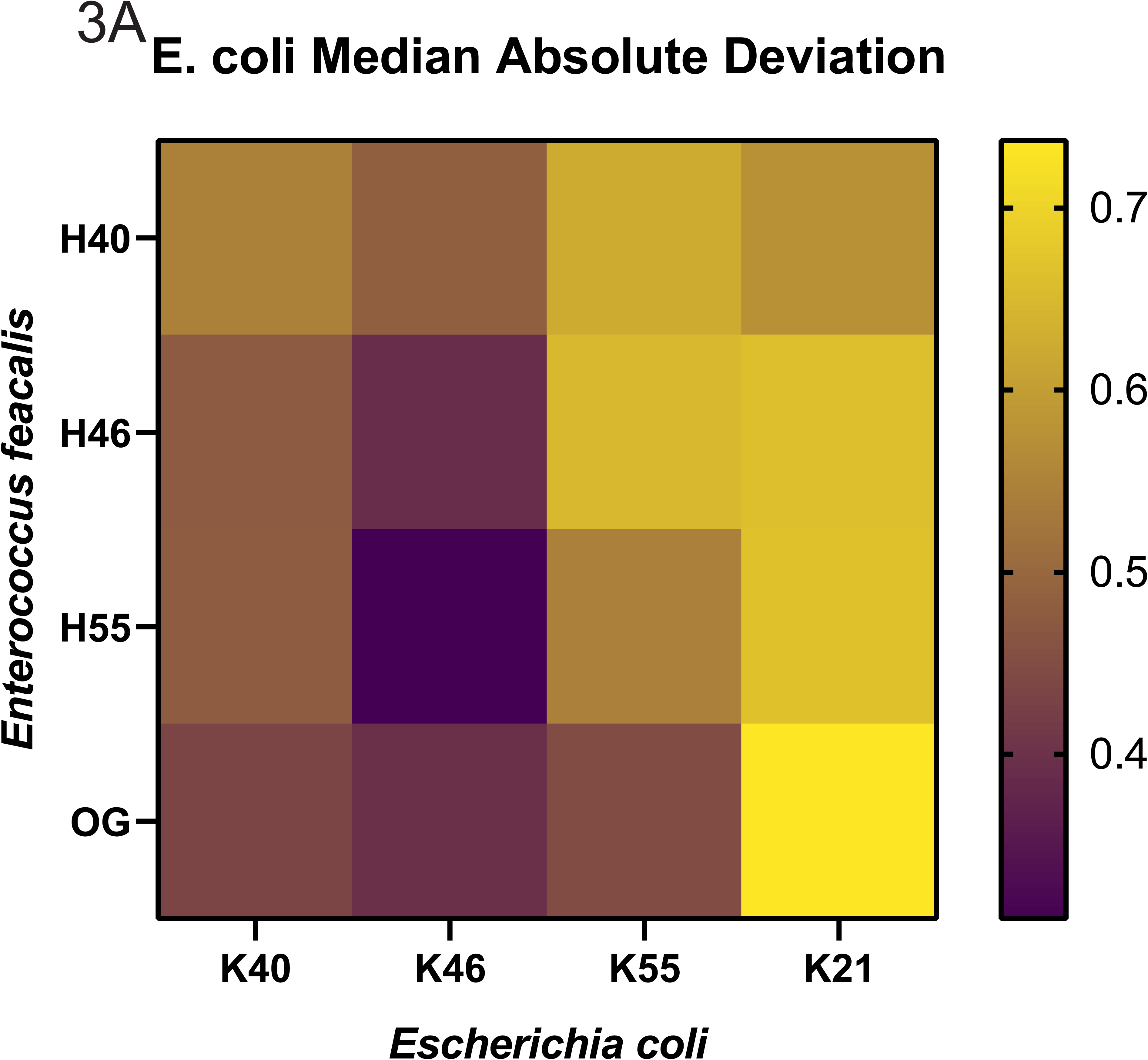
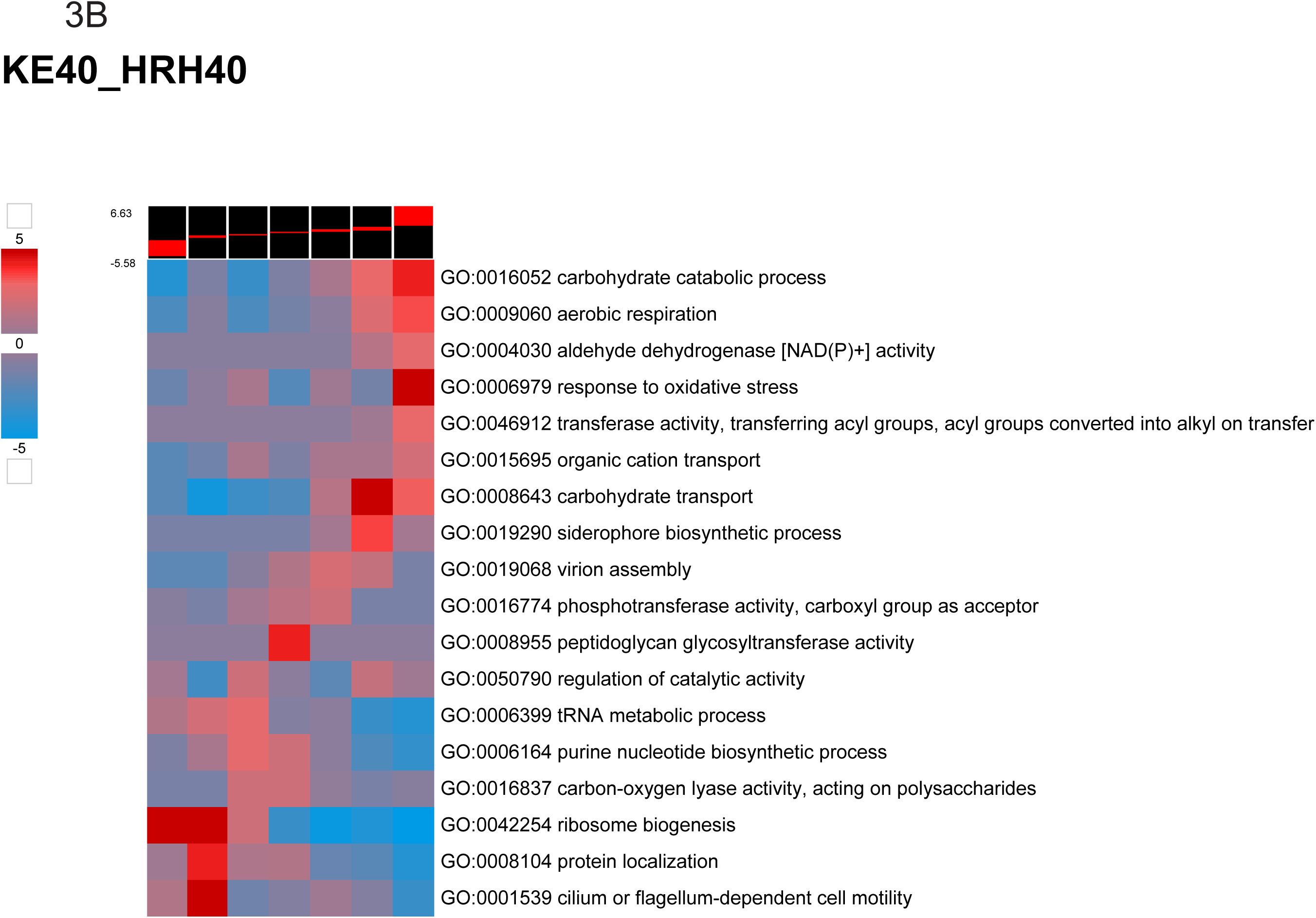
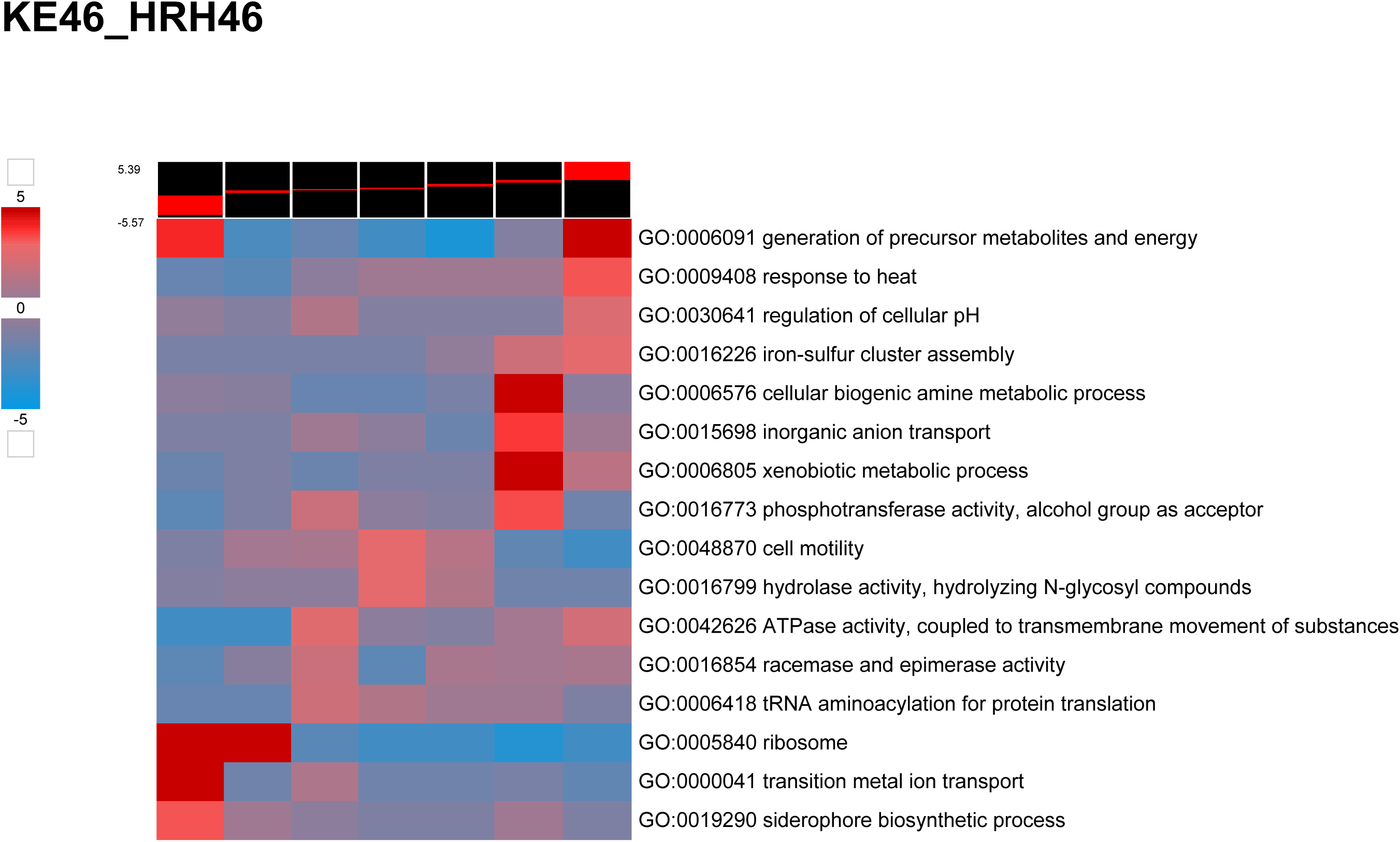
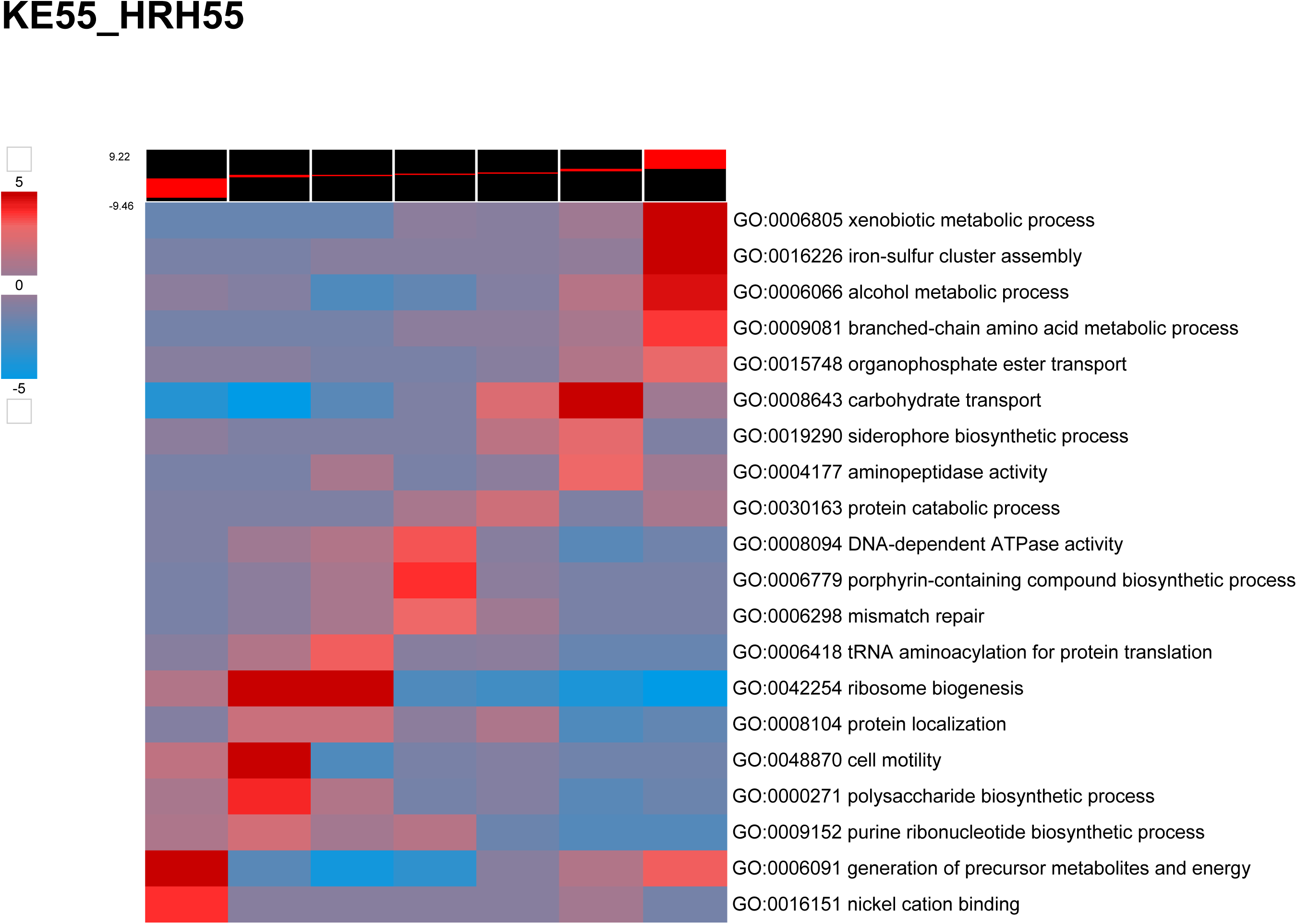
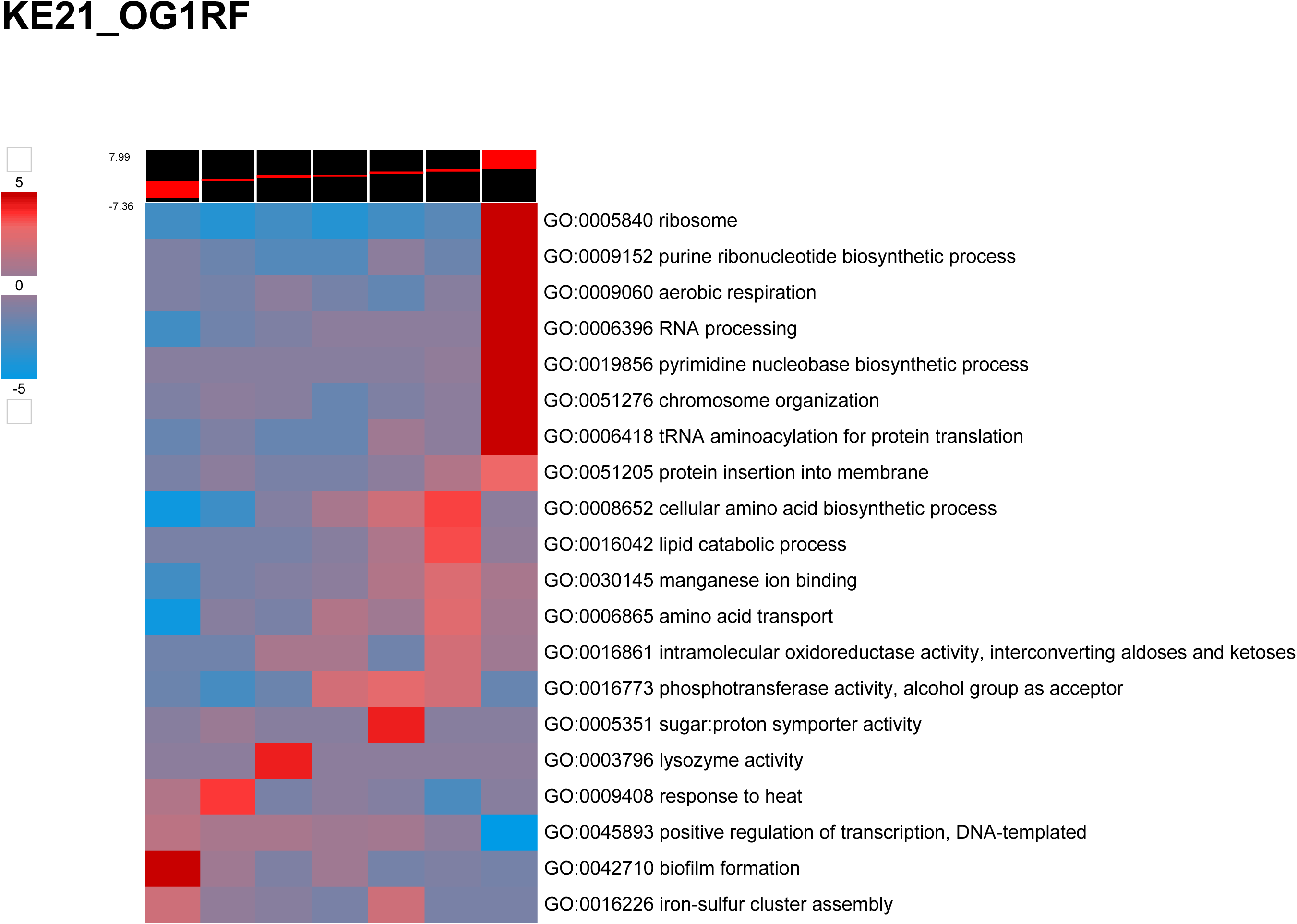
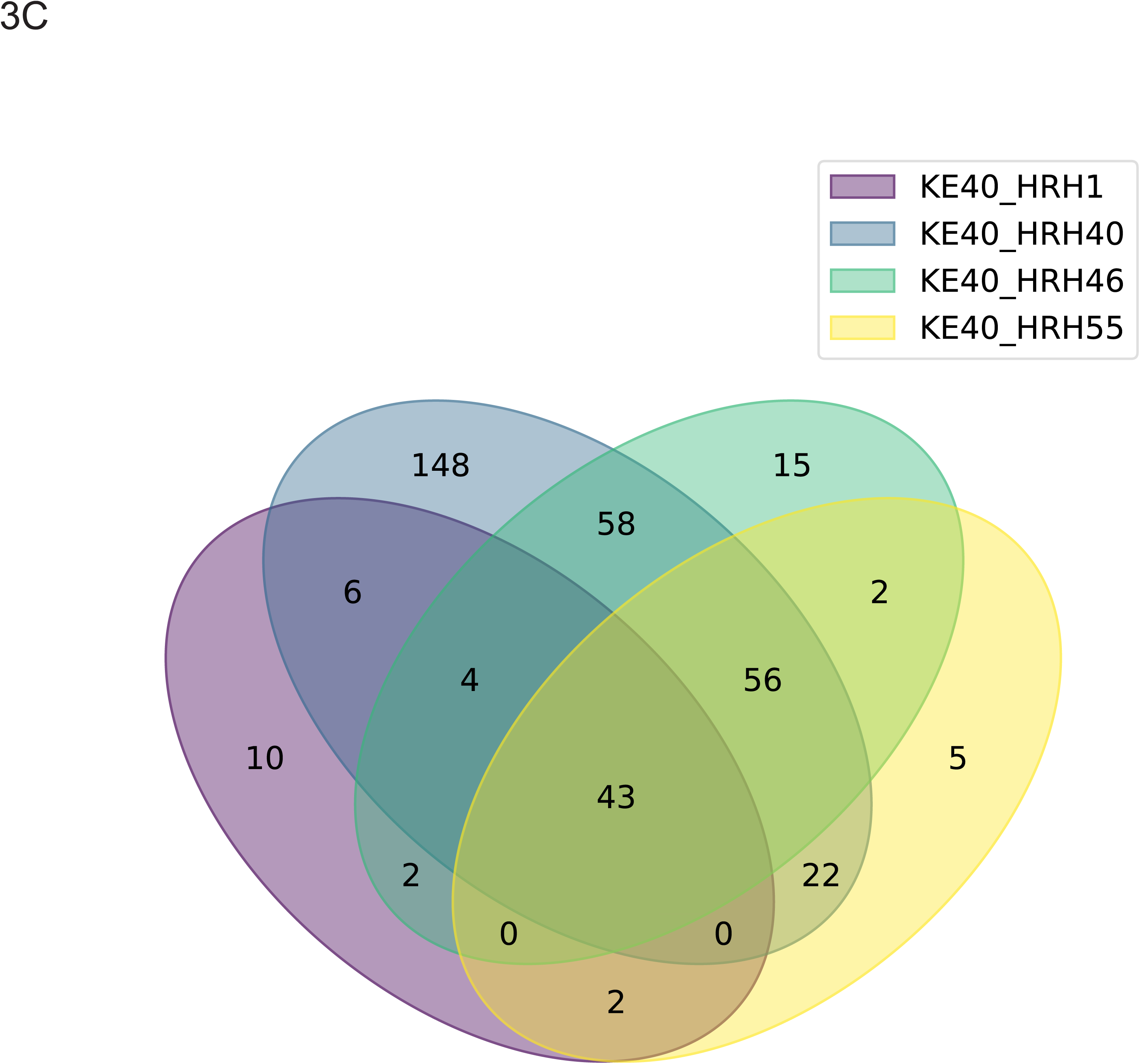
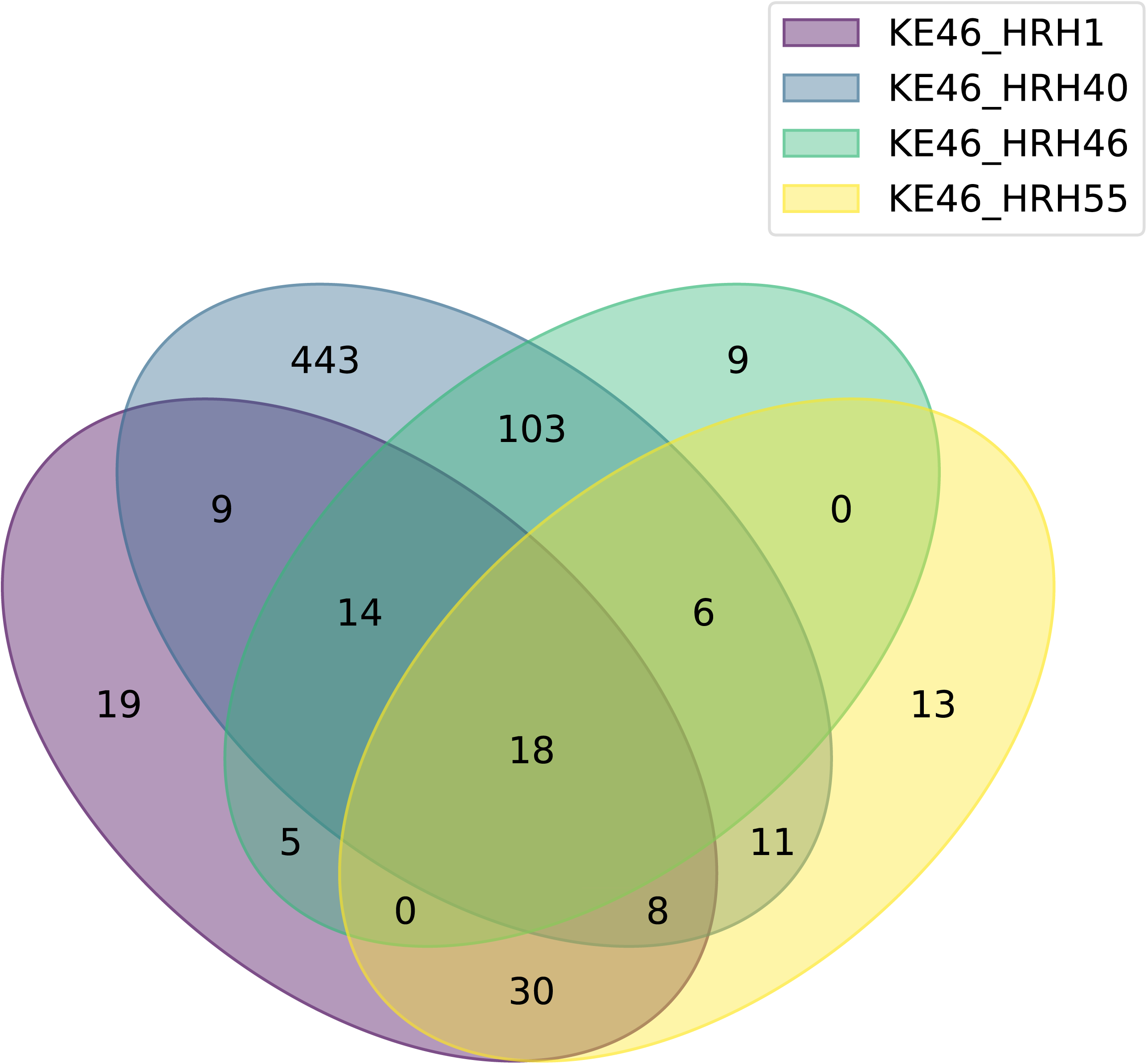
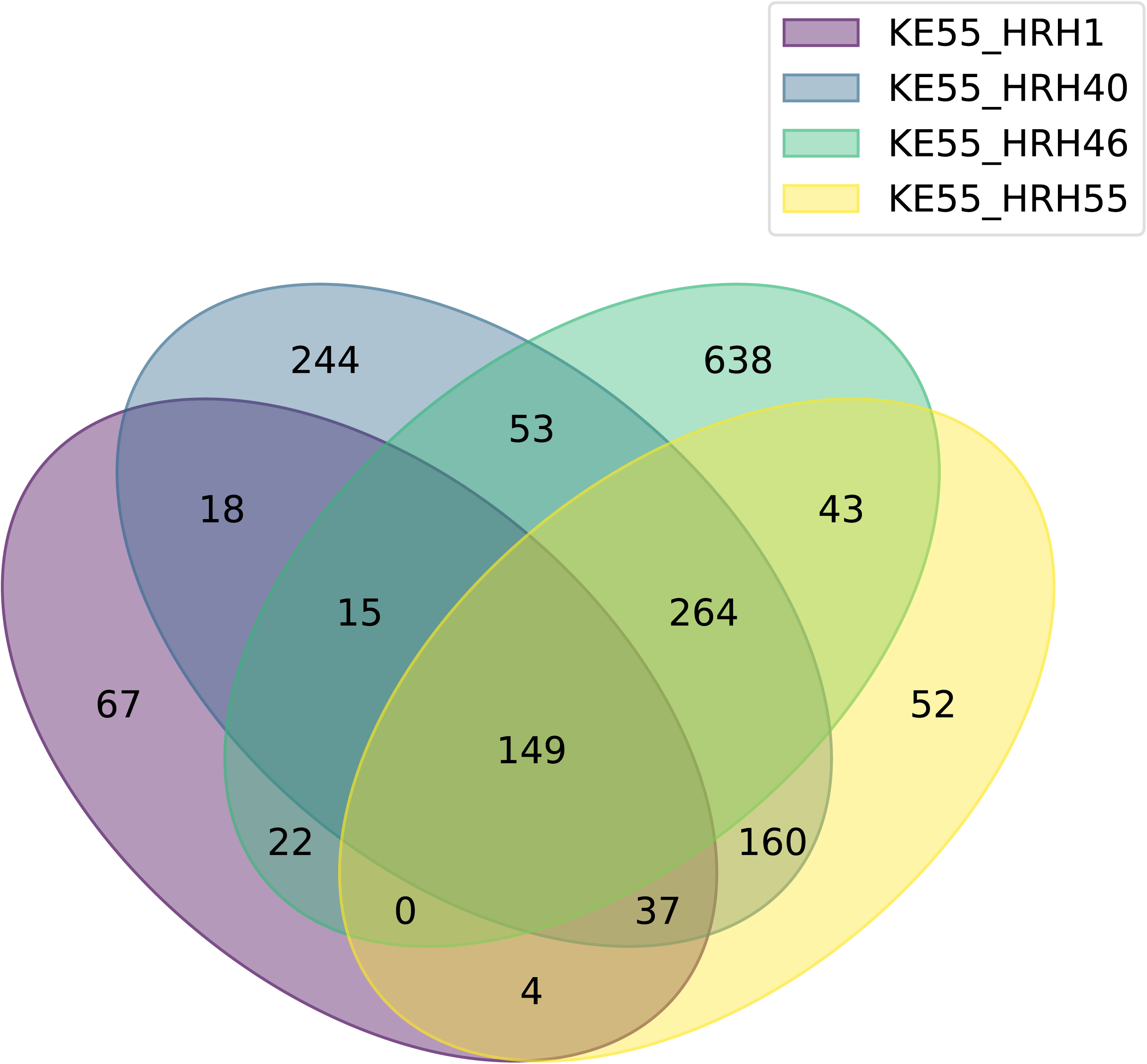
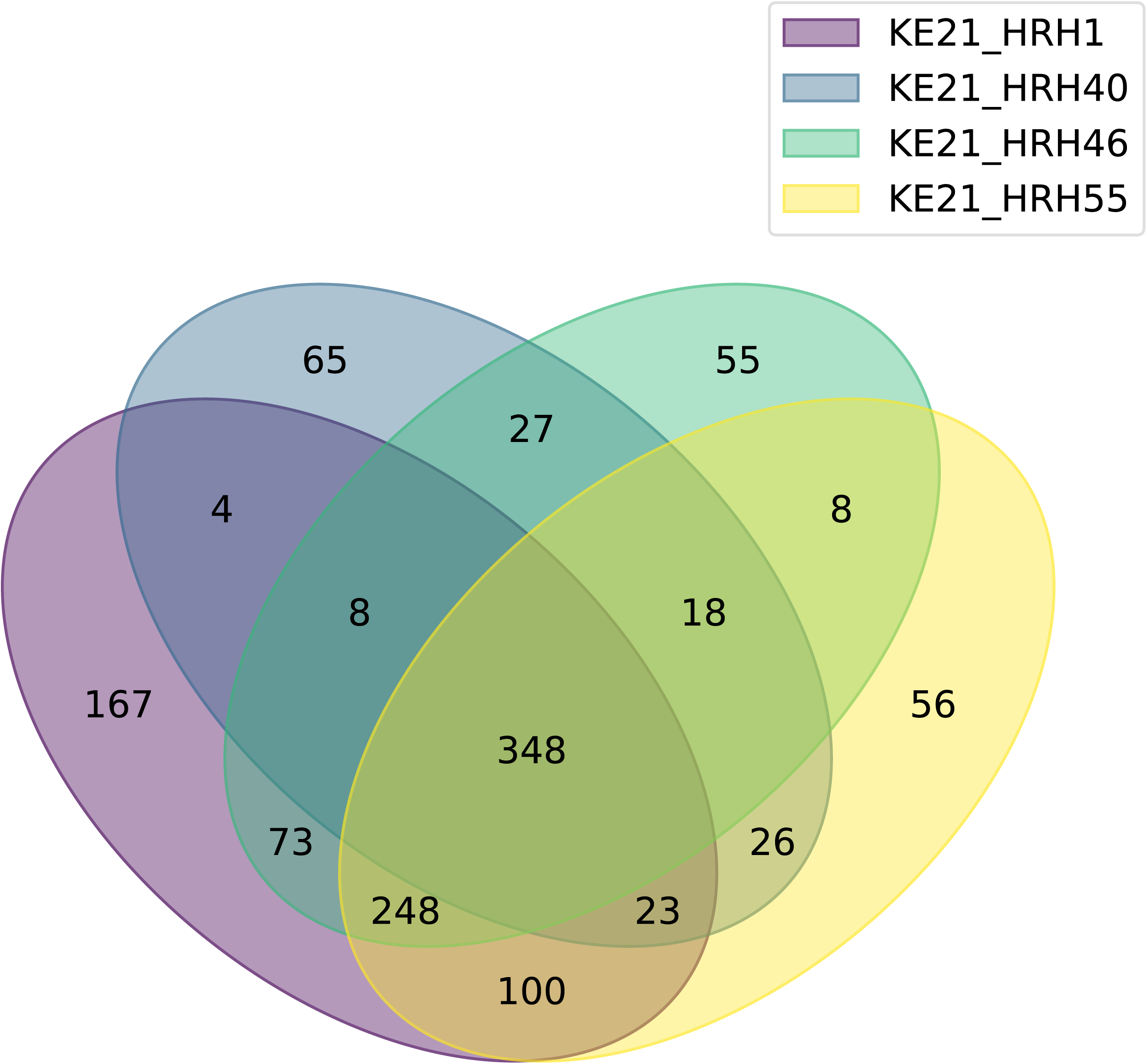
Functional Summary. A) Median absolute deviation of the Log2 Fold change of the Ec genes. Greater values reflect greater changes in the transcriptome. B) Heatmaps showing the changes in GO terms for the Ec genes. C) Venn diagrams showing the number of genes that significantly change in each pairing with an Ec strain.

The following sections describe transcript changes for individual genes by category. The heatmaps are ordered by the degree of enrichment in the most-upregulated bins, with two clear patterns emerging. First, Ef affected the non-B2 strains differently than the B2 strain KE21. Second, the effects of the different Ef strains, including the lab strain OG1RF, could not be distinguished which implies non-specific responses. The descriptions that follow apply to transcripts changes for most strain combinations.

### Ef had little effect on non-B2 strain core processes for the non-B2 Ec strains

For the non-B2 strains, the four Ef strains resulted in changes for only one category of genes, those coding for a few nucleoid-associated proteins. Increased transcripts were observed for Dps and reduced transcripts for Fis and IhfA (Fig 4A). Dps is associated with stationary phase and acid stress; Fis is associated with rapid growth; and IhfA is a component of a nucleoid-associated protein. Ef had no effect or modestly reduced transcripts from the core process genes of DNA replication, ribosomal proteins, RNA polymerase core subunits, and peptidoglycan synthesis and maturation (Fig 4 B-E). The one exception was *rpoS* transcripts which were higher for most Ef-non-B2 combinations (Fig 4D).

**Fig 4:**
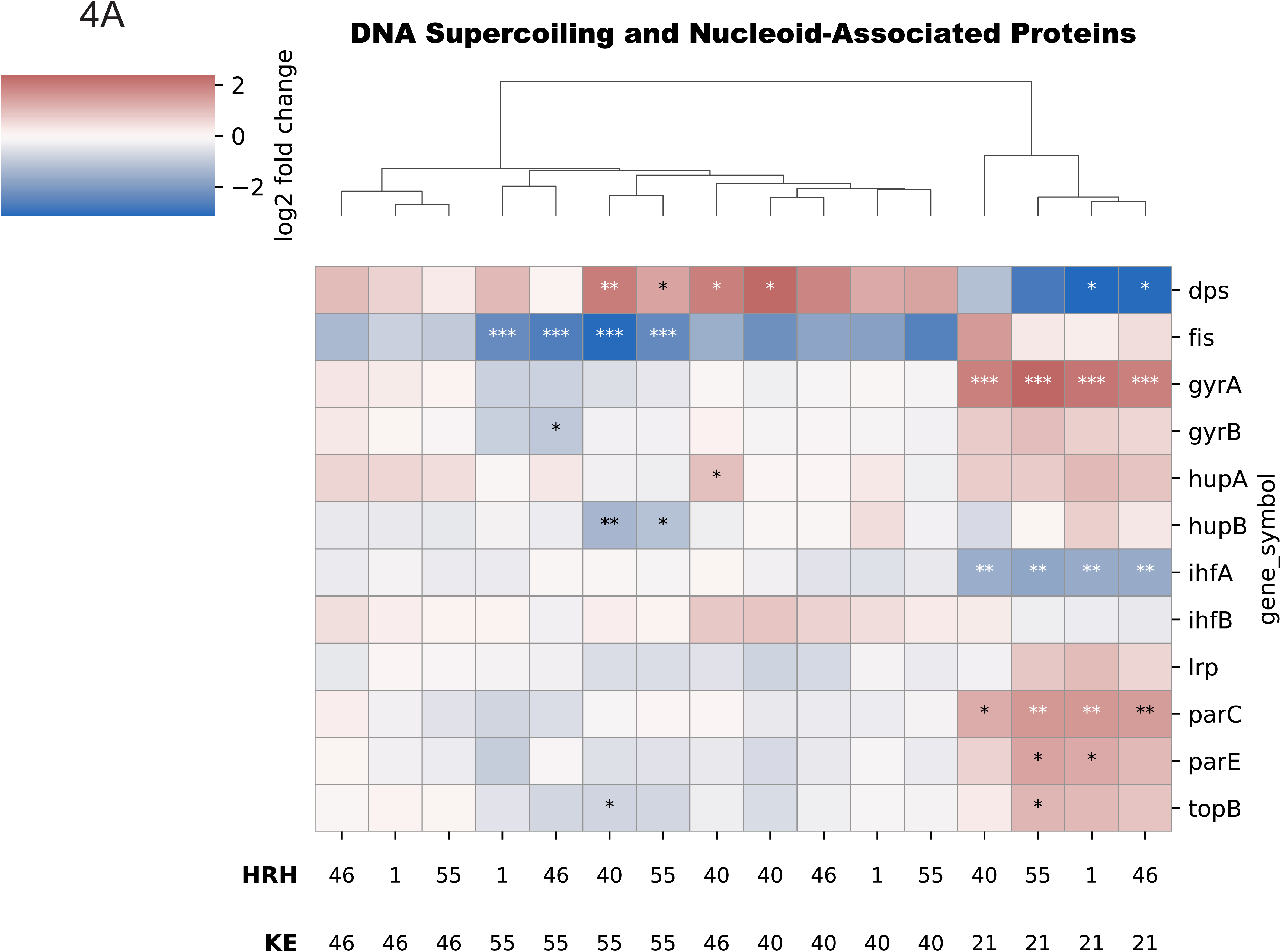
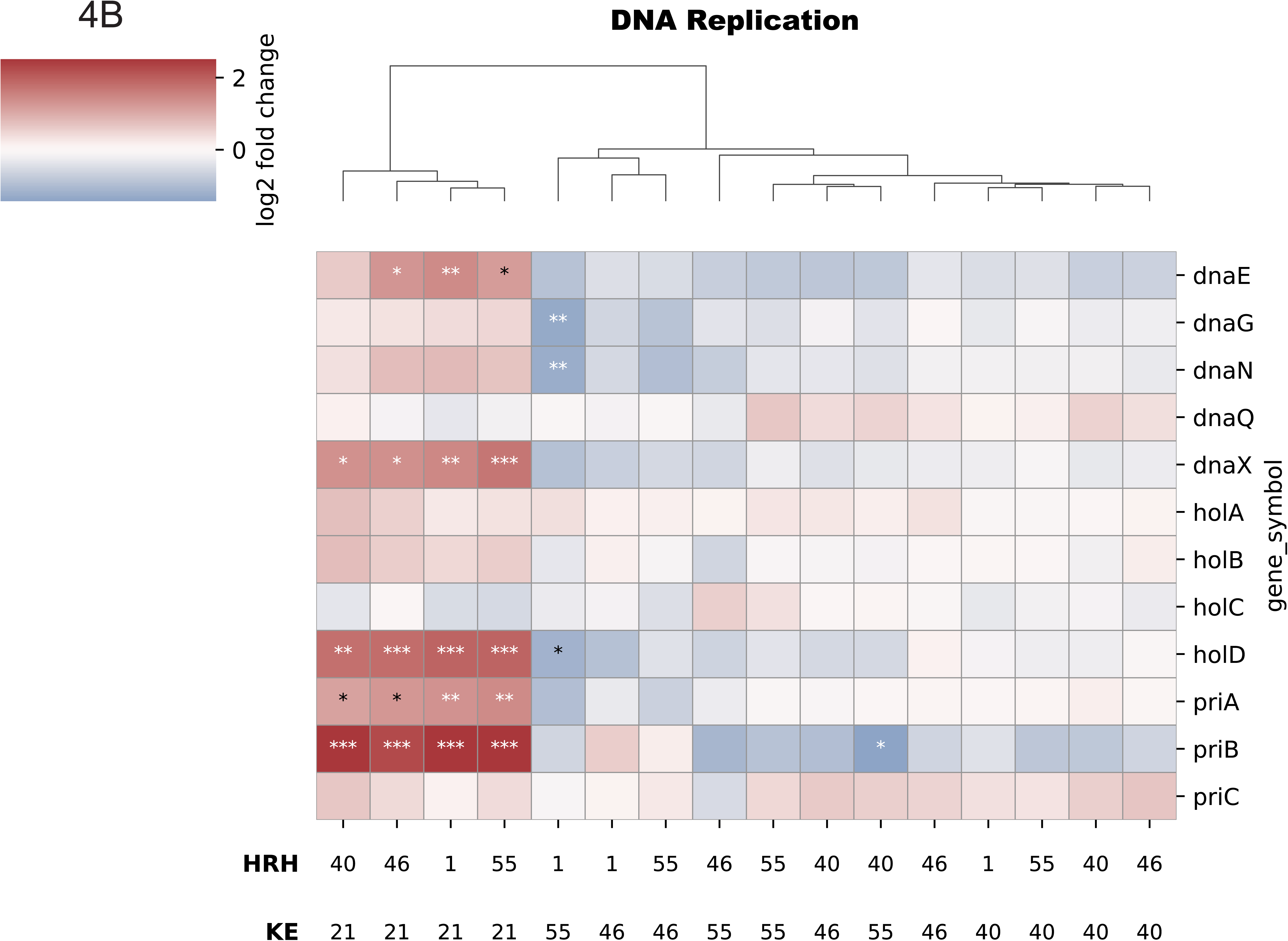
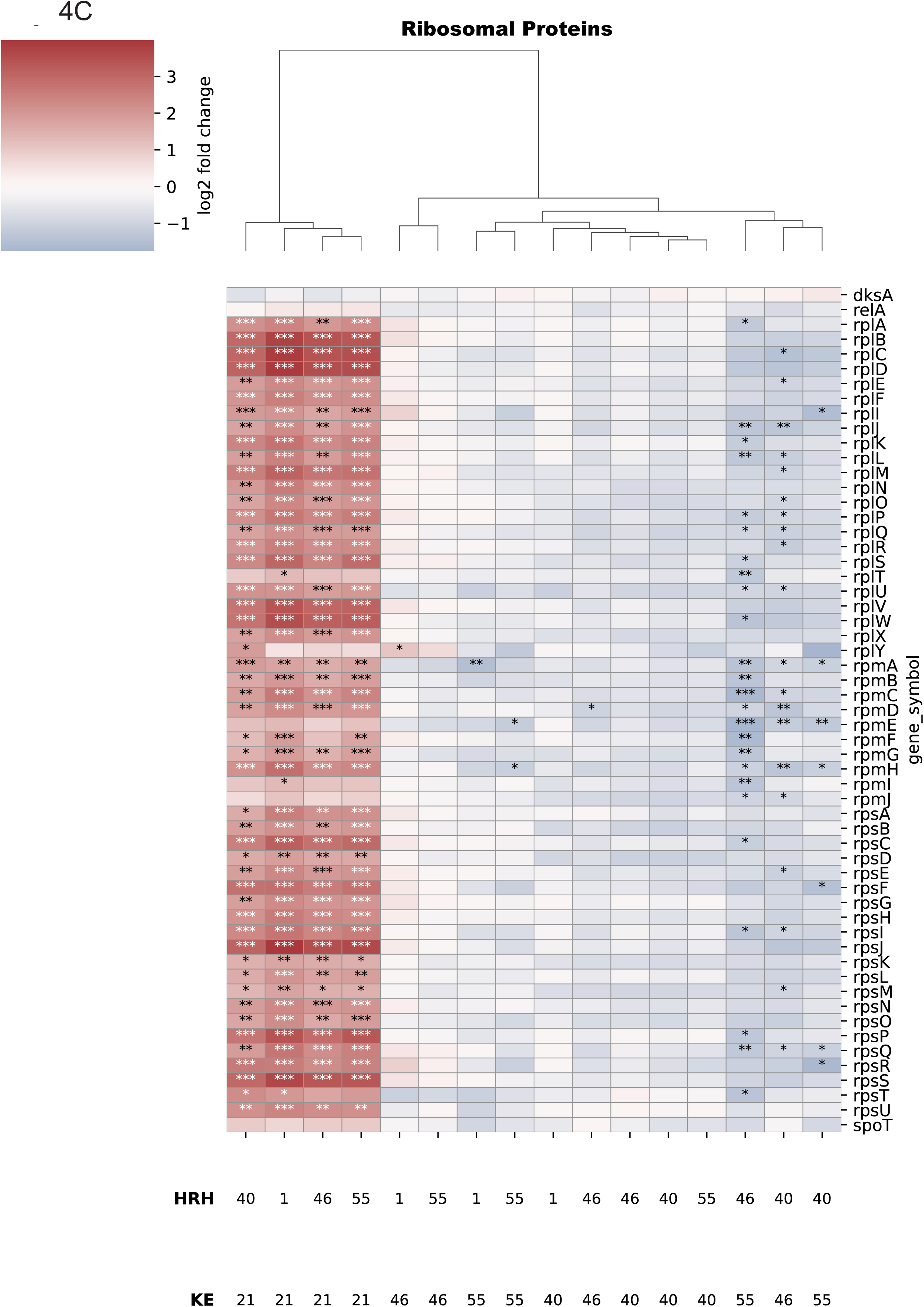
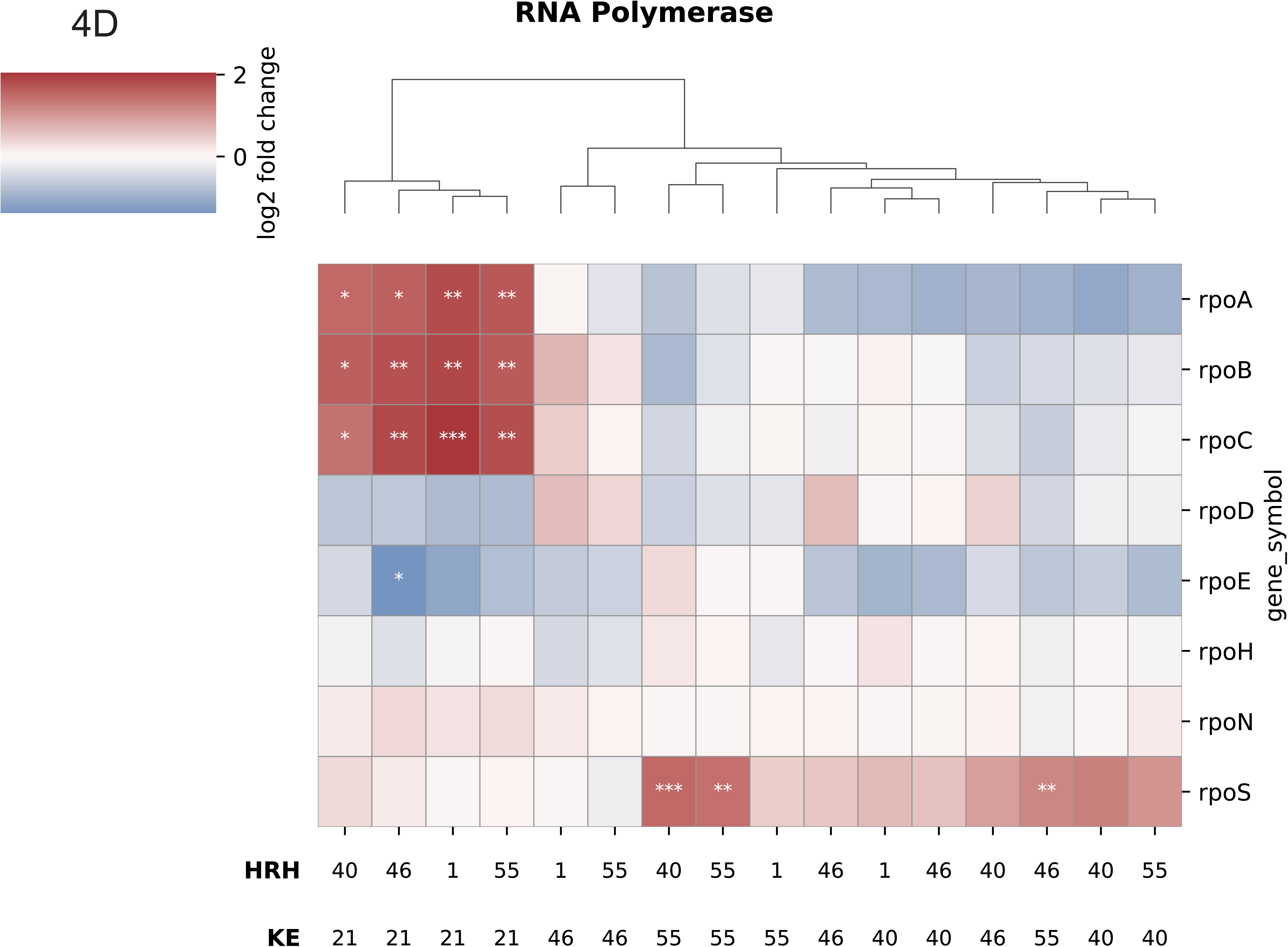
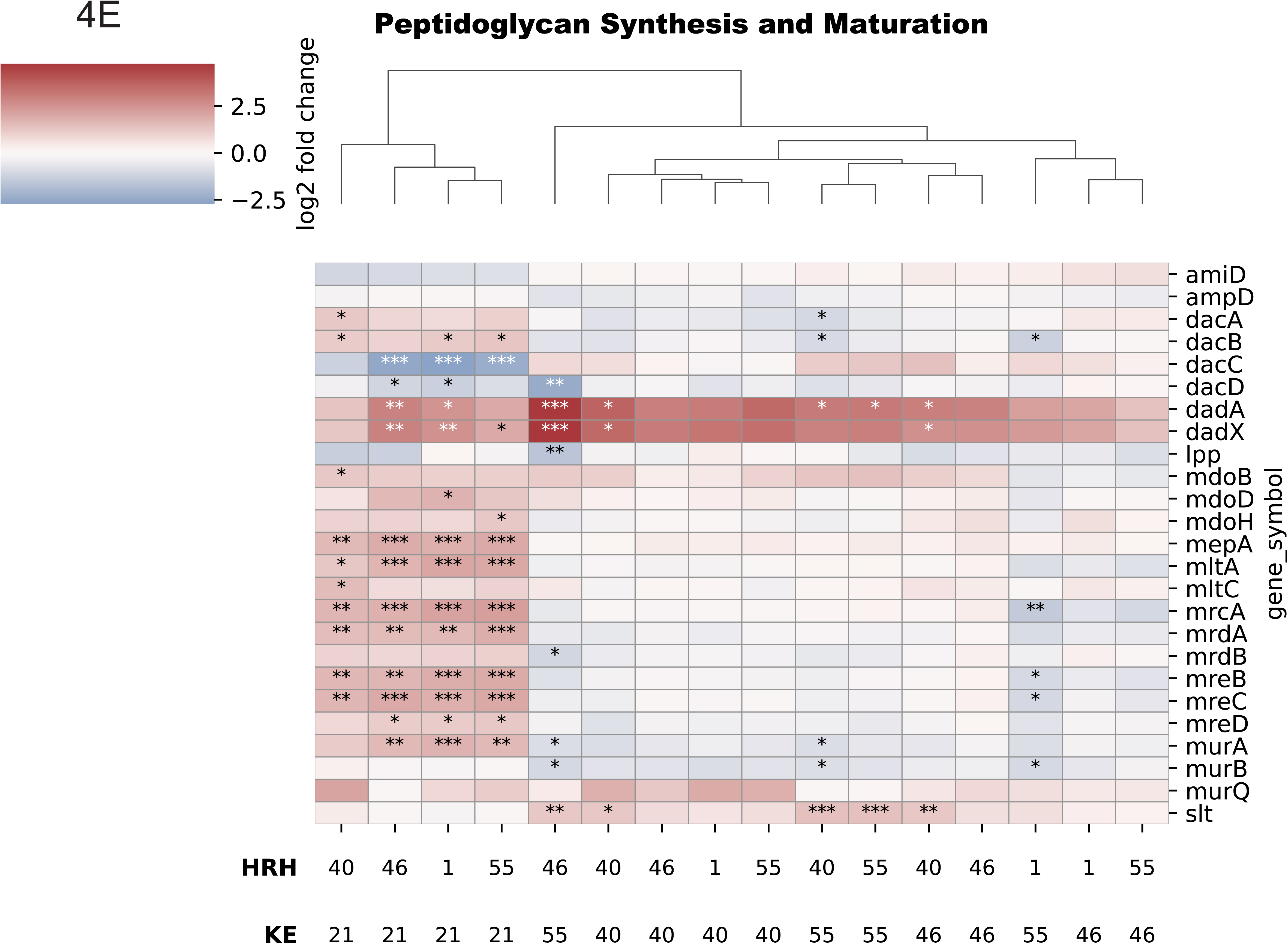
Transcript Changes for Ec Core Processes. Blue depicts decreases, red depicts increases in gene expression in a co-culture compared to the respective Ec monoculture. Stars represent q-values * <0.05 **<0.01 ***<0.001. A) DNA Supercoiling and Nucleoid Associated Proteins B) DNA Replication C) Ribosomal Proteins D) RNA Polymerase E) Peptidoglycan Biosynthesis and Maturation

### Ef had a large effect on the B2 Ec strain core processes

For the B2 strain control, the four Ef strains increased transcripts from the DNA gyrase subunit gene *gyrA* and the decantenating topoisomerase genes *parA*, *parC*, and *topB* (Fig 4A); DNA replication genes *dnaE*, *dnaX*, *holD*, *priA*, and *priB* (Fig 4B); ribosomal protein genes (Fig 4C); RNA polymerase core subunit genes *rpoA*, *rpoB*, and *rpoC* (Fig 4D); and peptidoglycan synthesis and maturation genes (Fig 4E). These transcript differences were not reflected in growth rate differences: for example, transcripts for ribosomal proteins are often four-fold higher with Ef, but growth is not four-fold faster.

### Ef effect on non-B2 strain metabolism

In contrast to effects on core processes, Ef affected non-B2 Ec strain transcripts from numerous metabolic genes. Ef co-culture induced Ec transcripts for all TCA cycle enzymes except for the pyruvate dehydrogenase subunits AceE, AceF, and Lpd (Fig 5A). In addition, Ef induced transcripts for genes coding for MaeB (the malic enzyme), LldD (quinone-dependent L-lactate dehydrogenase), PoxB (quinone-dependent pyruvate oxidase), Acs (acetyl-CoA synthetase), and the glyoxylate shunt enzymes AceA (isocitrate lyase), AceB (malate synthase), and AceK (a kinase that phosphorylates and inactivates isocitrate dehydrogenase) (Fig 5A).

**Fig 5:**
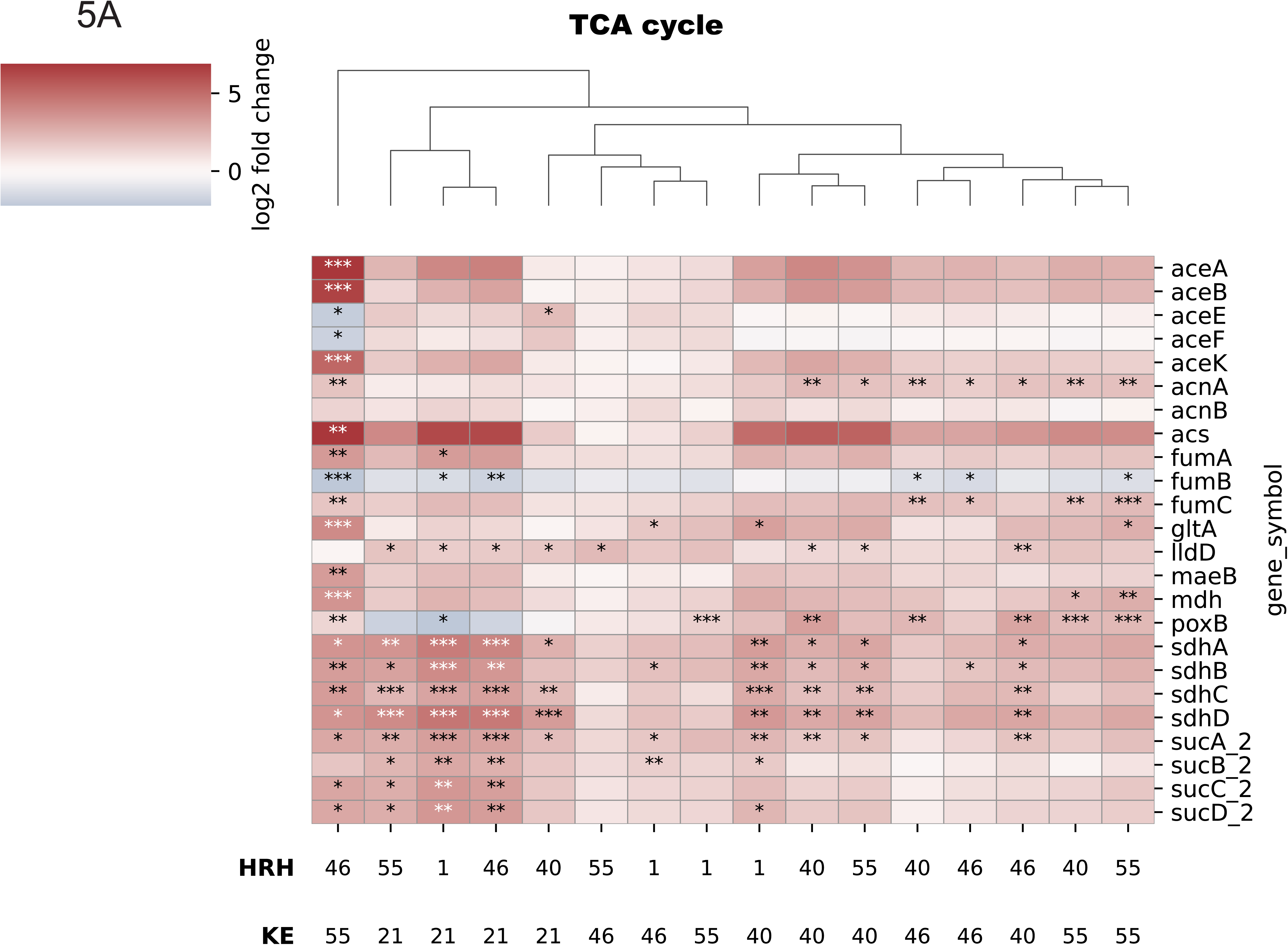
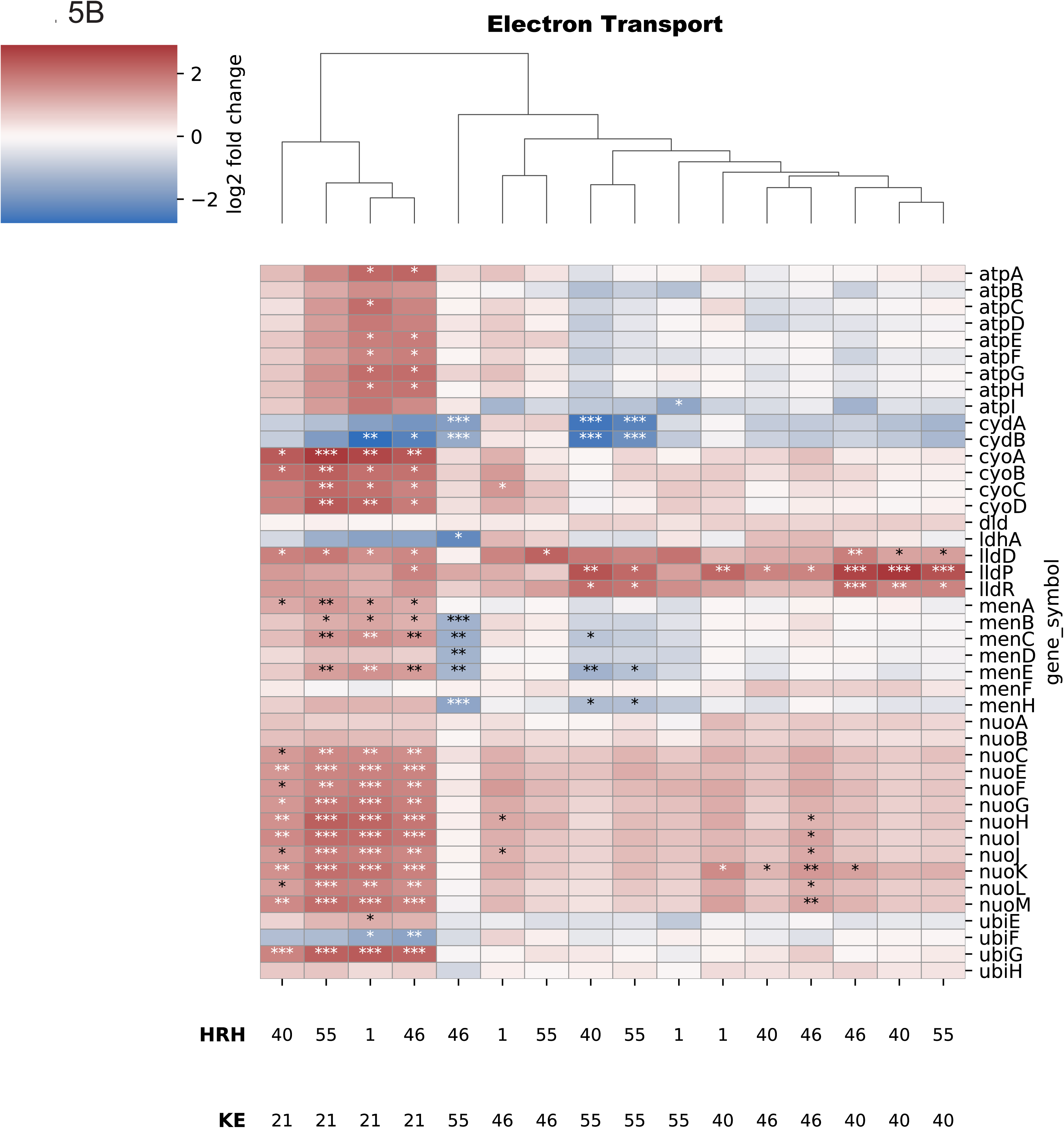
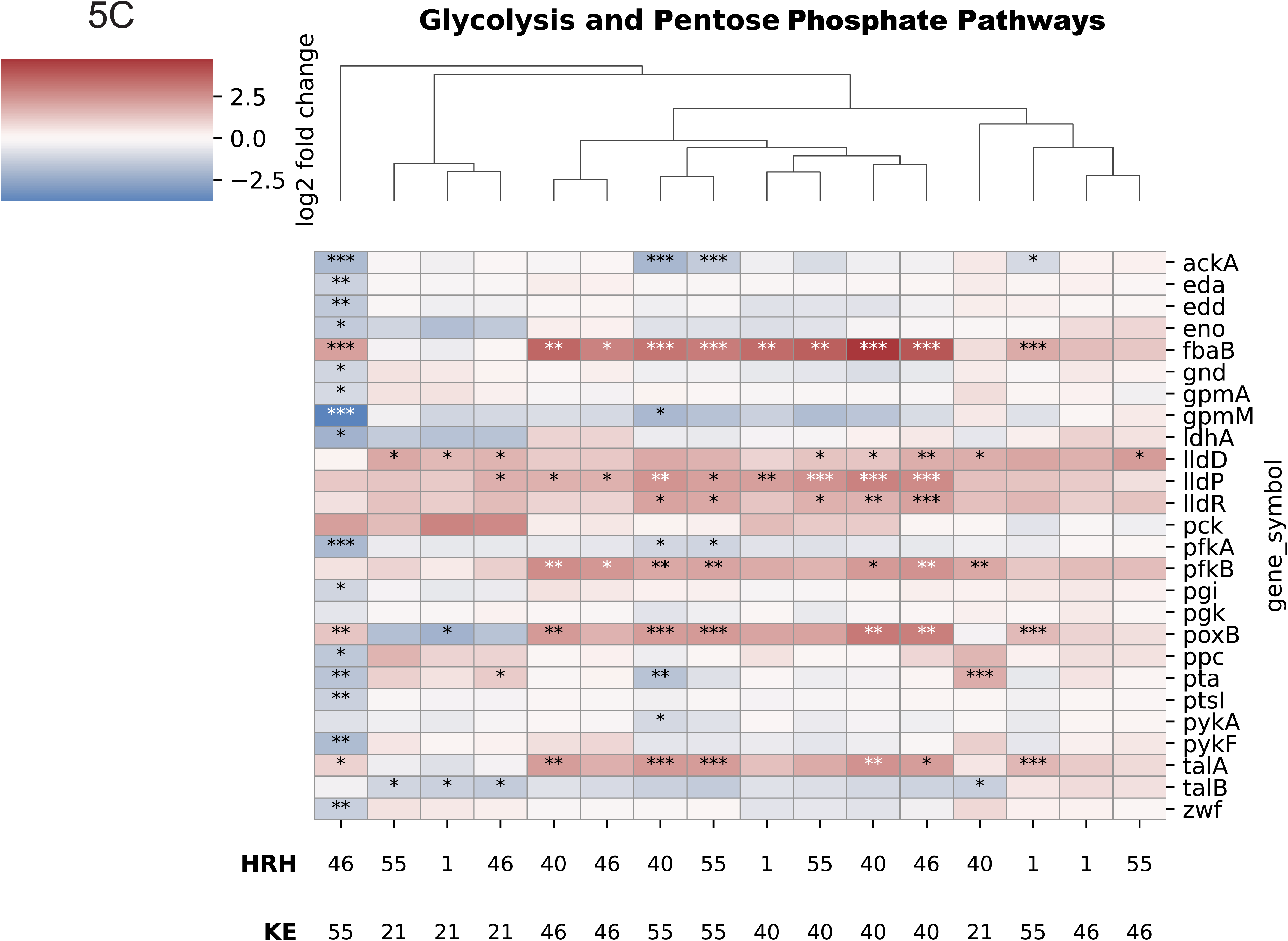
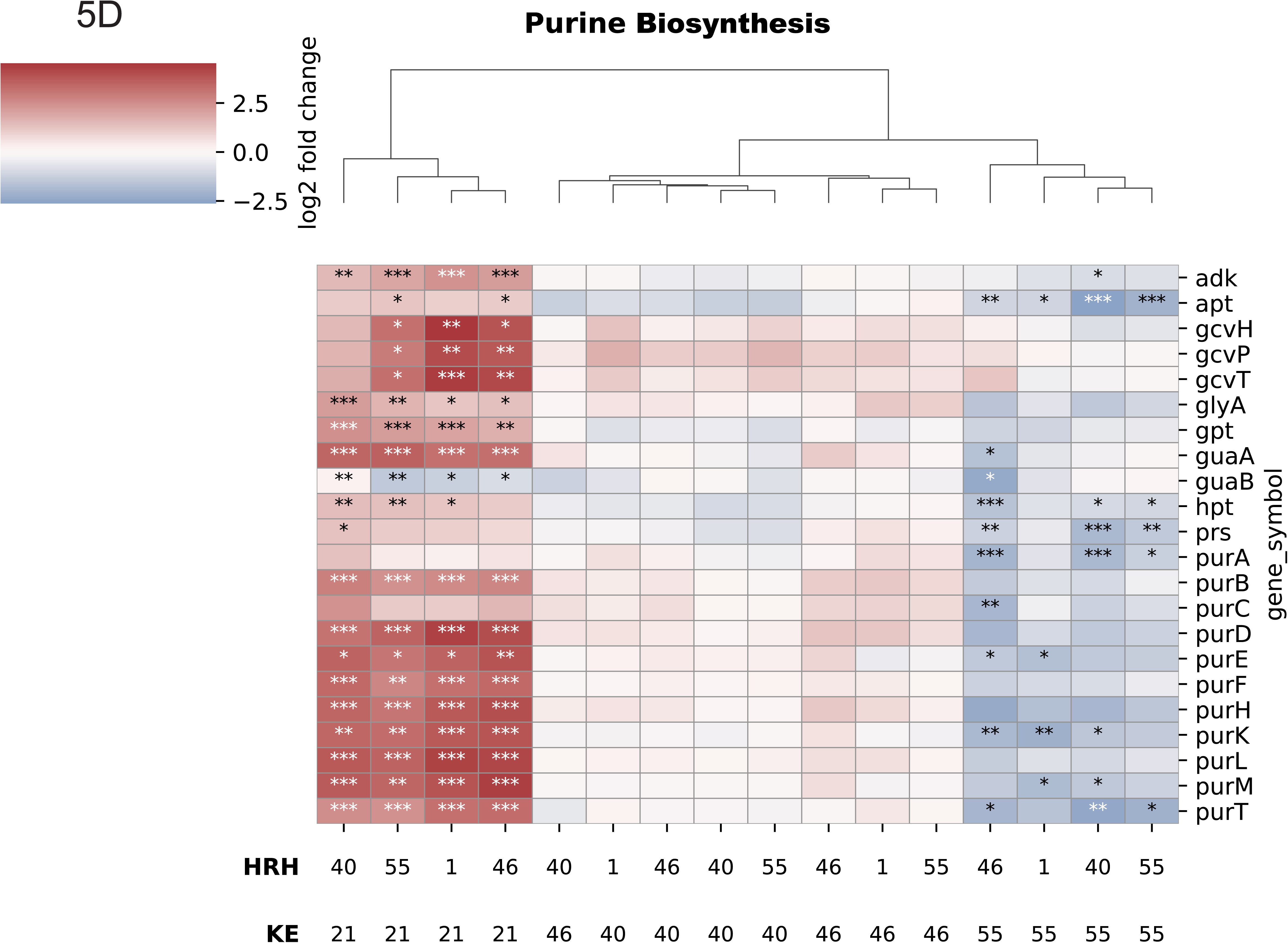
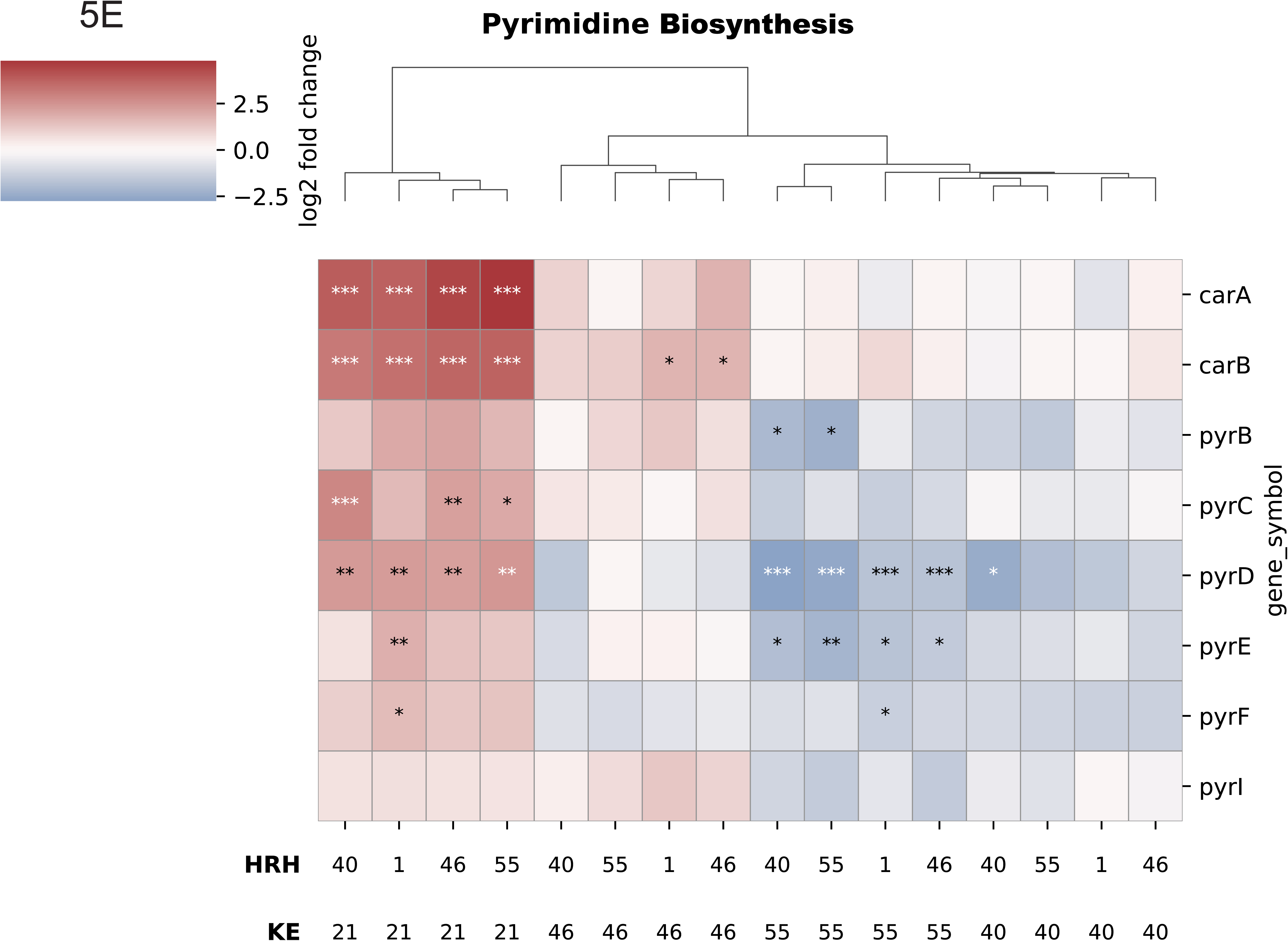
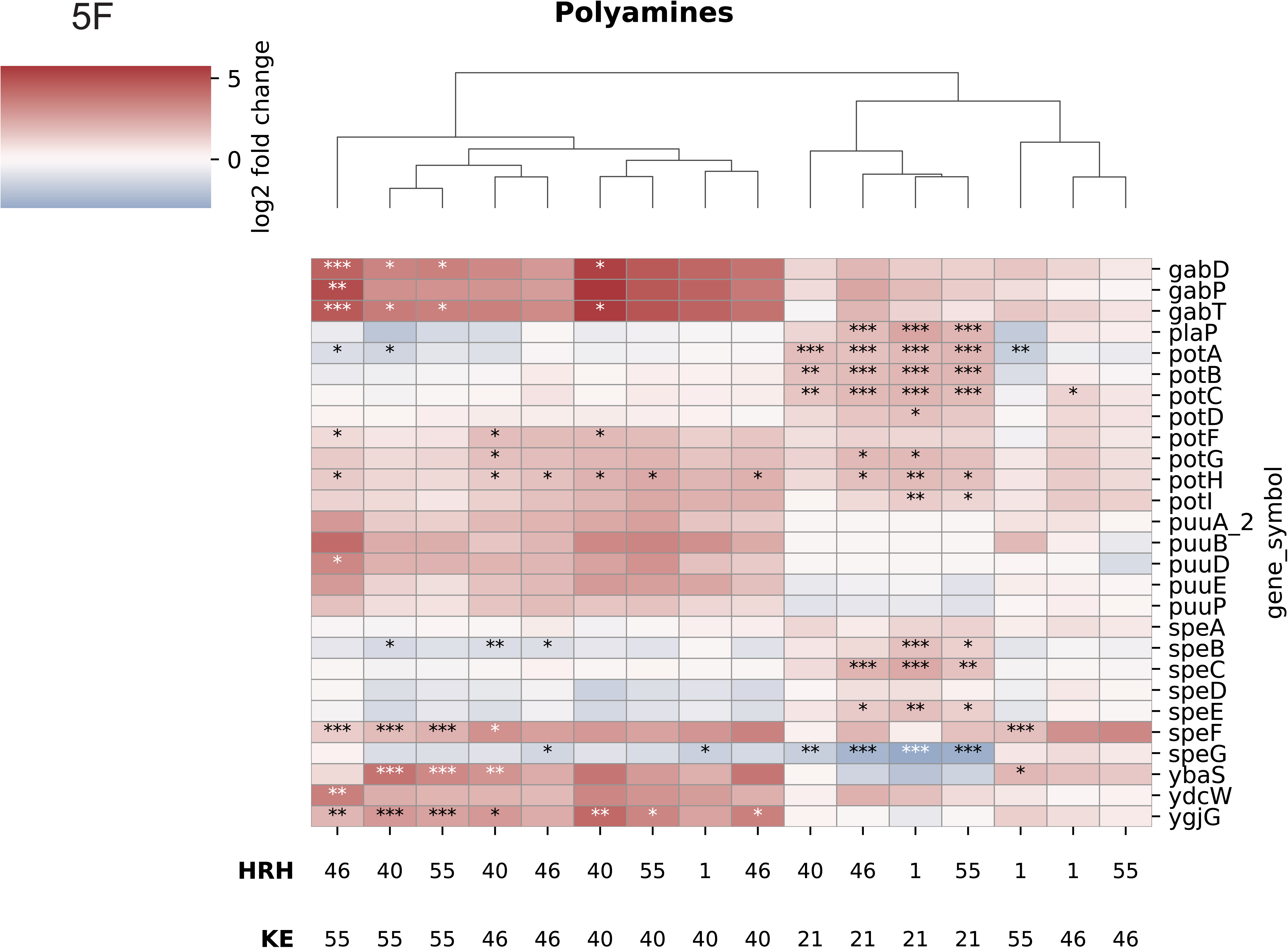
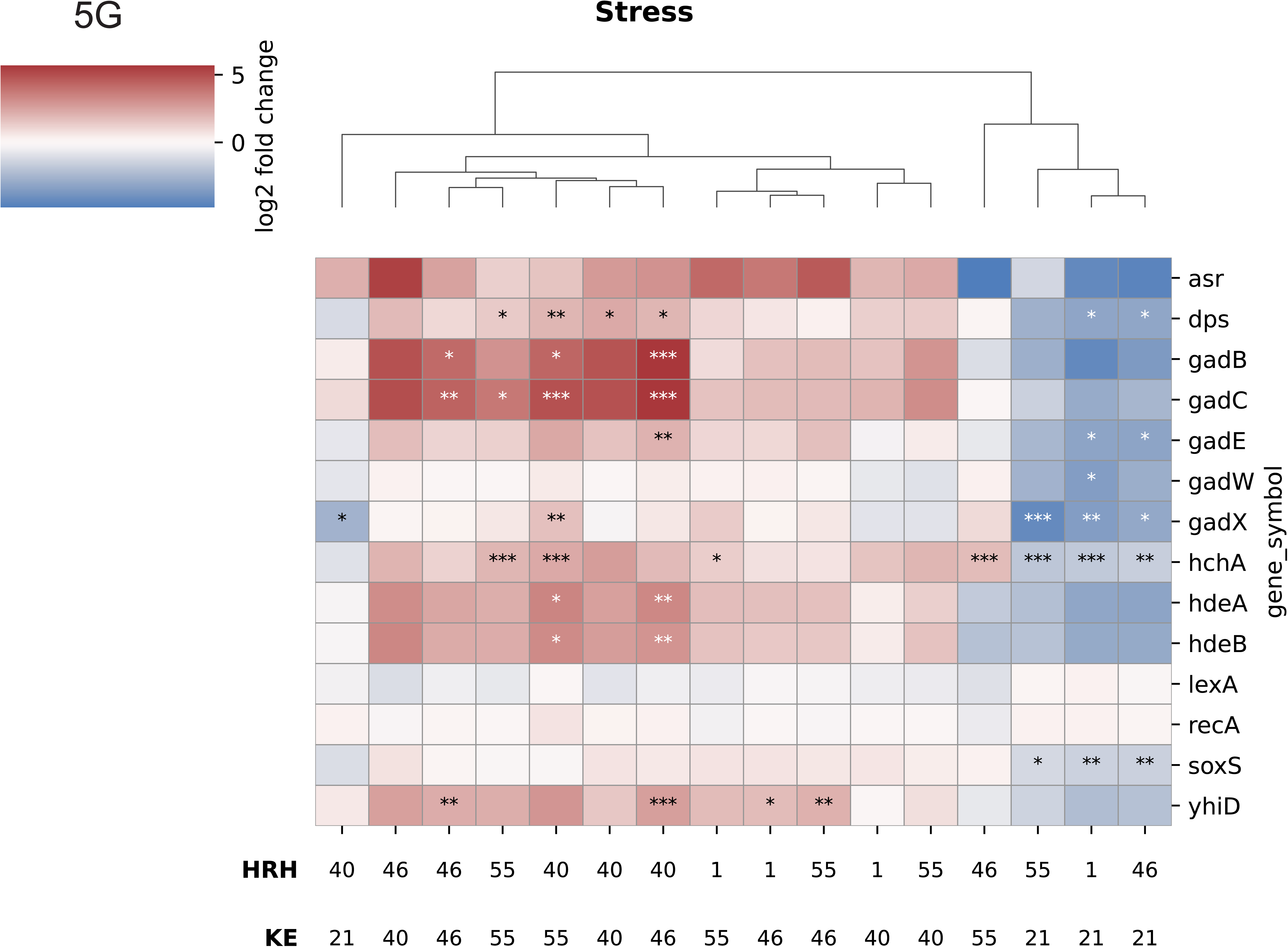
Transcript Changes for Ec Metabolic Genes. Blue depicts decreases, red depicts increases in gene expression in a co-culture compared to the respective Ec monoculture. Stars represent q-values * <0.05 **<0.01 ***<0.001. A) TCA Cycle B) Electron Transport C) Glycolysis and Pentose Phosphate Pathway D) Purine Biosynthesis E) Pyrimidine Biosynthesis F) Polyamines G) Stress

Ef induced more transcripts for energy-conserving NADH dehydrogenase 1 (*nuo* operon gene products), but fewer transcripts for the non-energy-conserving high-oxygen-affinity cytochrome oxidase *bd* (CydA-CydB) (Fig 5B). Transcripts for other electron transport components were largely unaffected (Fig 5B).

Transcripts for a few enzymes of glycolysis and the non-oxidative pentose phosphate pathway were also induced: phosphofructokinase 2 (PfkB), a fructose-bisphosphate aldolase isozyme (FbaB), glyceraldehyde-3-phosphate dehydrogenase (GapA), transaldolase A (TalA), and transketolase 2 (TktB) (Fig 5C).

Ef had little effect on transcripts from purine and pyrimidine synthesis pathways for KE40 and KE46 but generally reduced these transcripts for KE55 (Fig 5D and E).

Ef co-culture generally increased transcripts from several genes of polyamine metabolism (Fig 5F). These included genes for putrescine synthesis from ornithine (*speF*), putrescine transport (*potFGHI*), putrescine catabolism (*patA*, *patD*, *puuE*, and *puuP*), and γ-aminobutyrate (GABA) catabolism (*gabDPT*). GABA is a product of putrescine catabolism.

Ef co-culturing resulted in more transcripts for several low pH-induced genes (Fig 5G): *asr* (acid shock), *dps* (DNA protection), *gadB* (proton-consuming glutamate decarboxylase), *hchA* (an acid stress chaperone), the *hdeAB* operon (periplasmic acid stress chaperones), and *speF* (ornithine decarboxylase). Co-culturing generally did not affect transcripts for the oxidative stress and DNA damage genes *oxyR*, *soxS*, *lexA*, and *recA* which suggests the relative absence of these stresses. The exception was HRH40 which induced transcripts for these four genes when co-cultured with KE21 (B2), KE46 (B1), and KE55 (D).

### Differential Ef effect on metabolism of the group B2 strain

Like its effect on non-B2 strains, Ef increased transcripts in the B2 strain for TCA cycle and glyoxylate shunt enzymes (Fig 5A). Unlike the response of non-B2 strains, Ef modestly increased B2 strain transcripts for the pyruvate dehydrogenase subunits AceE and AceF (Fig 5A), greatly increased transcripts for NADH dehydrogenase 1 (the *nuo* operon) (Fig 5B), and increased transcripts for enzymes that synthesize menaquinone, cytochrome *bo* oxidase (for a high oxygen environment), and the ATP synthase complex (the *atp* operon) (Fig 5B). In addition to induced transcripts for genes of macromolecular synthesis and structure (Fig 4), Ef resulted in more transcripts for all enzymes of purine and pyrimidine metabolism, including GlnA (purine synthesis consumes glutamine), and GlyA and the GcvHTP complex (both generate a purine and pyrimidine intermediate) (Fig 5D and E). We conclude that Ef potentially stimulated energy metabolism to a greater extent in the B2 strain. Ef induced transcription from two polyamine transport systems (*potABCD* and *potFGHI*), only the latter was induced in the non-B2 strains, but did not induce the putrescine catabolic pathway genes (*patA*, *patD*, and *gabDTP*) (Fig 5F). B2 strains lack the major PuuA-initiated putrescine catabolic pathway: there are no transcripts for genes of the *puu* operons in the 26 B2 strains that we have examined [(24) and unpublished observation]. These results suggest that the B2 strain transports but does not degrade putrescine. Unlike the effect on the non-B2 strains, Ef co-culture reduced transcripts from acid stress genes and had no effect on other stress genes (Fig 5G).

### Differences between the non-B2 and B2 Ec strain responses of EF

For the non-B2 strains, Ef induced transcripts for metabolic enzymes that convert ornithine via putrescine and GABA to the TCA cycle intermediate succinate. Specific transcript changes are shown in Fig 5 and the deduced pathway changes are summarized in Fig 6. The major difference is that Ef induces transcripts in non-B2 strains for enzymes that degrade ornithine ― Ef produces ornithine (29) ― to TCA cycle intermediates for energy generation, while the B2 strain does not degrade ornithine past putrescine, i.e., putrescine is not an energy source.

**Fig 6:**
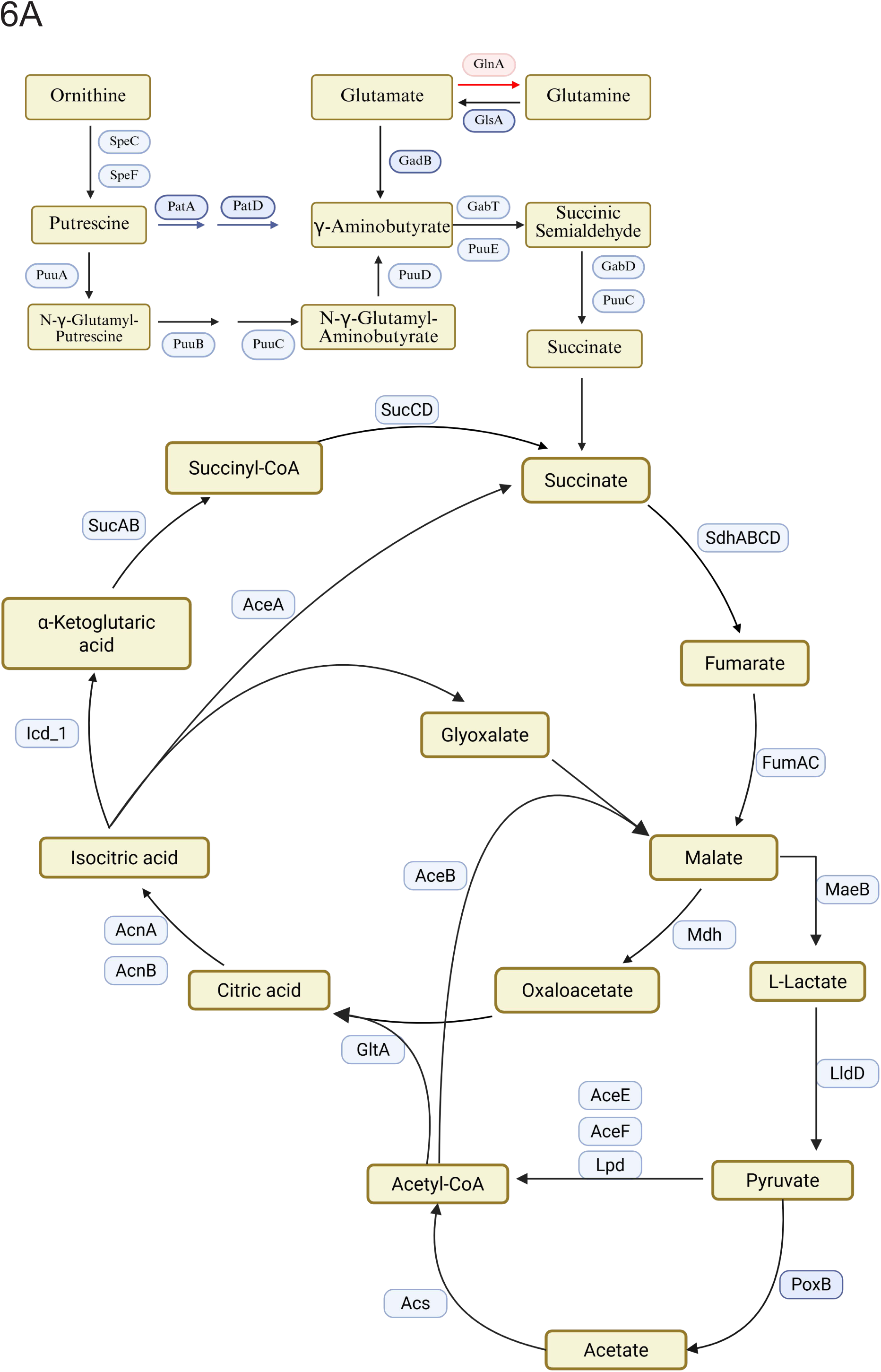
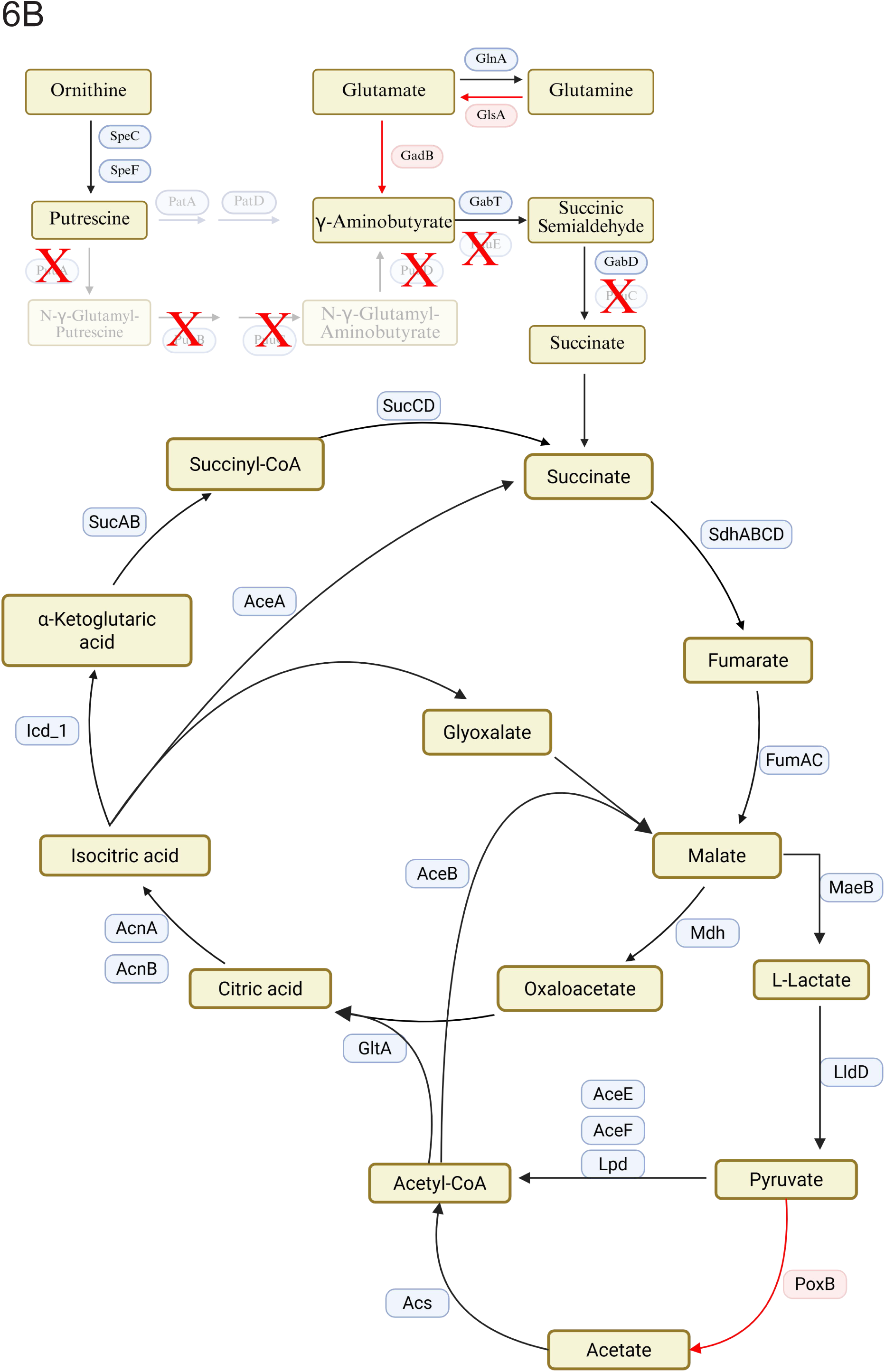
Deduced Pathway Changes. A) Gene expression changes in the co-cultures of Non-B2 strains of Ec compared to the monocultures. Increases are shown in black, decreased genes are shown in red. B) Gene expression changes in the co-cultures of the B2 strain of Ec compared to the monoculture. Increases are shown in black, decreased genes are shown in red, transparent genes are unchanged between monoculture and co-cultures, and transparent genes with X’s depict genes absent from the B2 genome.

### Ec induces two different Ef responses

HRH40 responded similarly to non-B2 and B2 Ec strains with fewer transcripts for genes of ribosomal proteins (Fig 7A), core RNA polymerase subunits (Fig 7B), and pyrimidine synthesis enzymes (Fig 7C); and increased transcripts for genes of purine synthesis enzymes (Fig 7D), and chaperones and proteases (Fig 7E). The discordance between purine and pyrimidine synthesis genes results from control of the purine genes by factors other than biosynthetic requirements (30). In summary, Ec strains non-specifically induce stress in HRH40.

**Fig 7:**
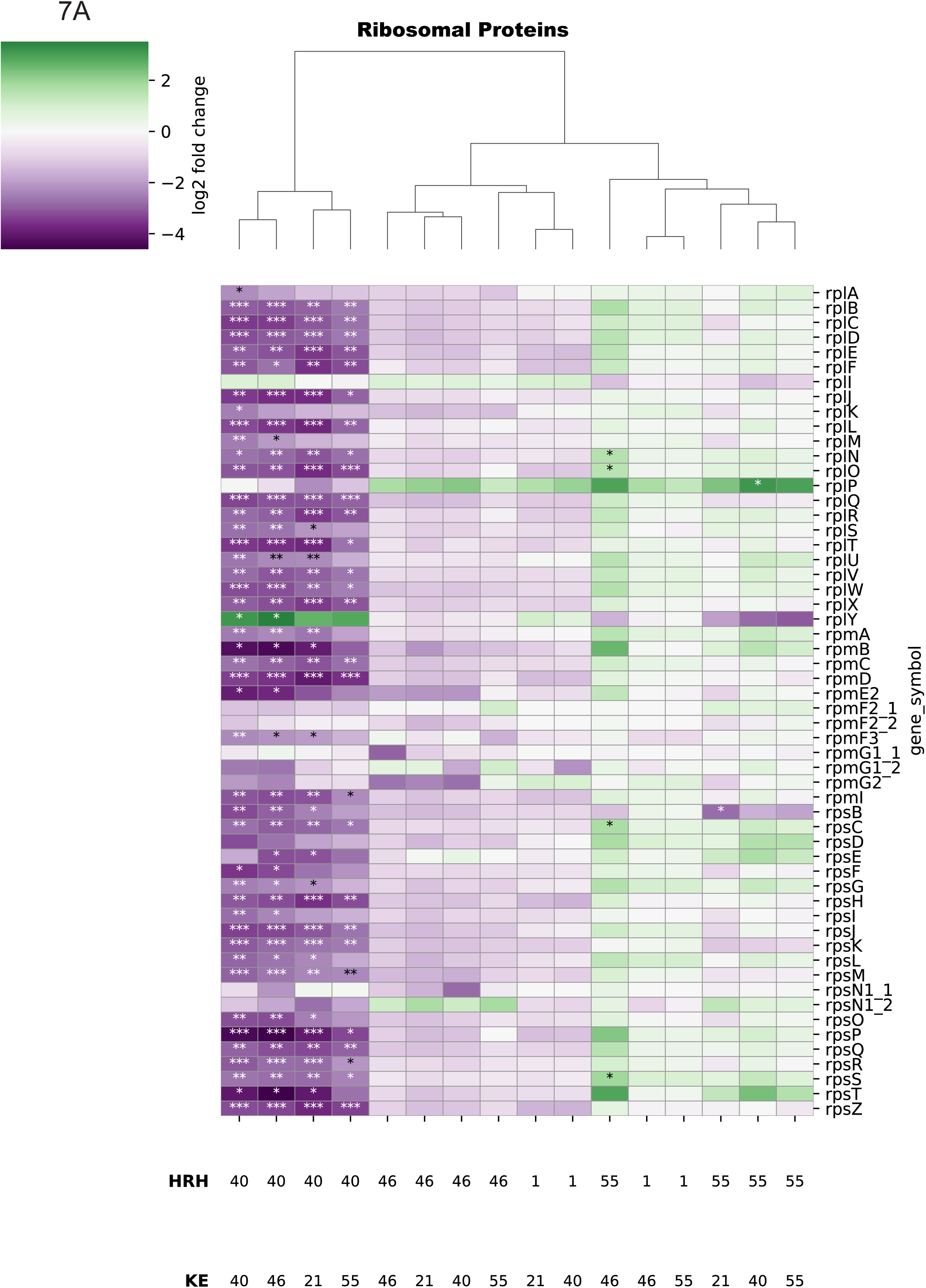
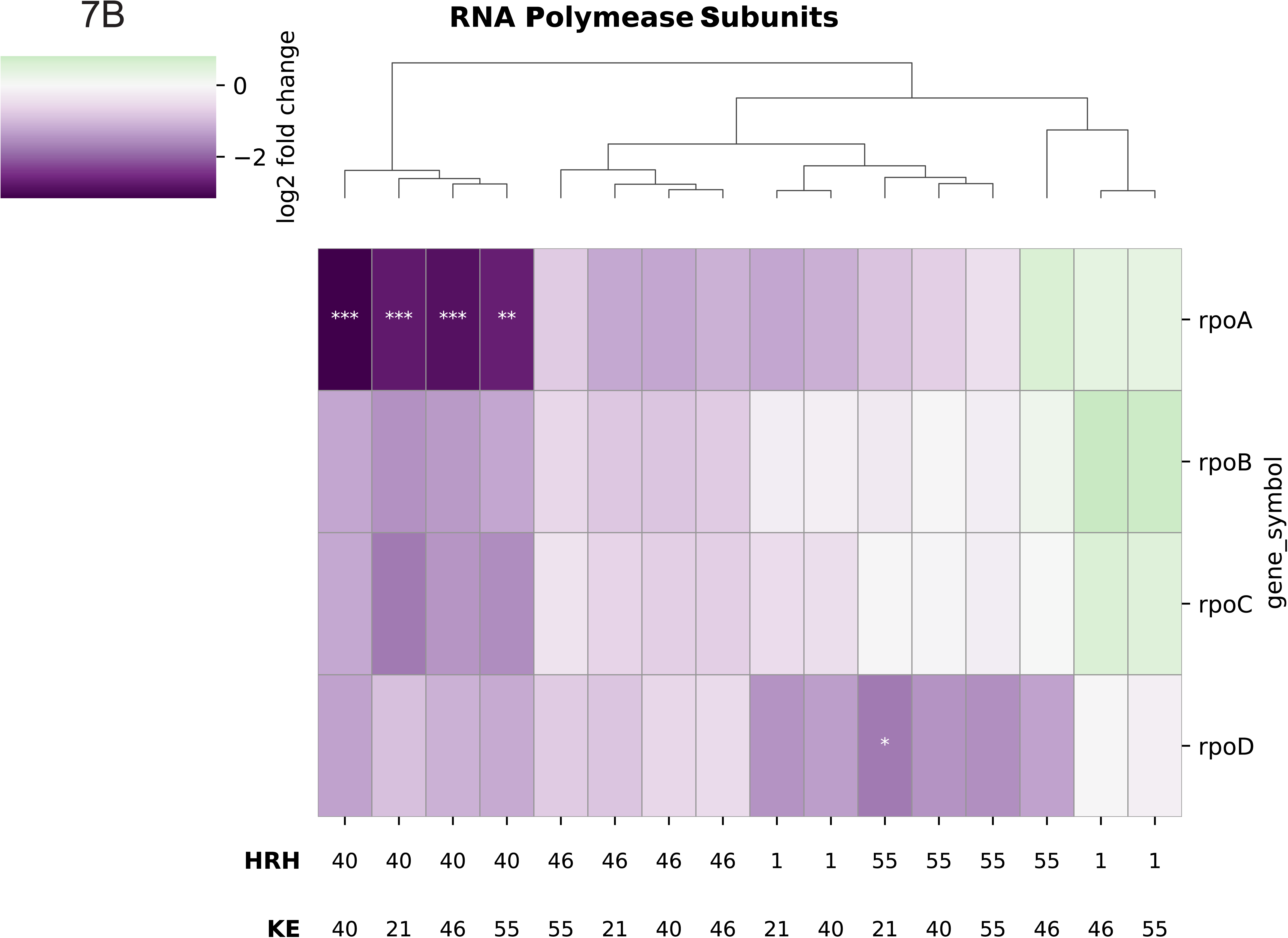
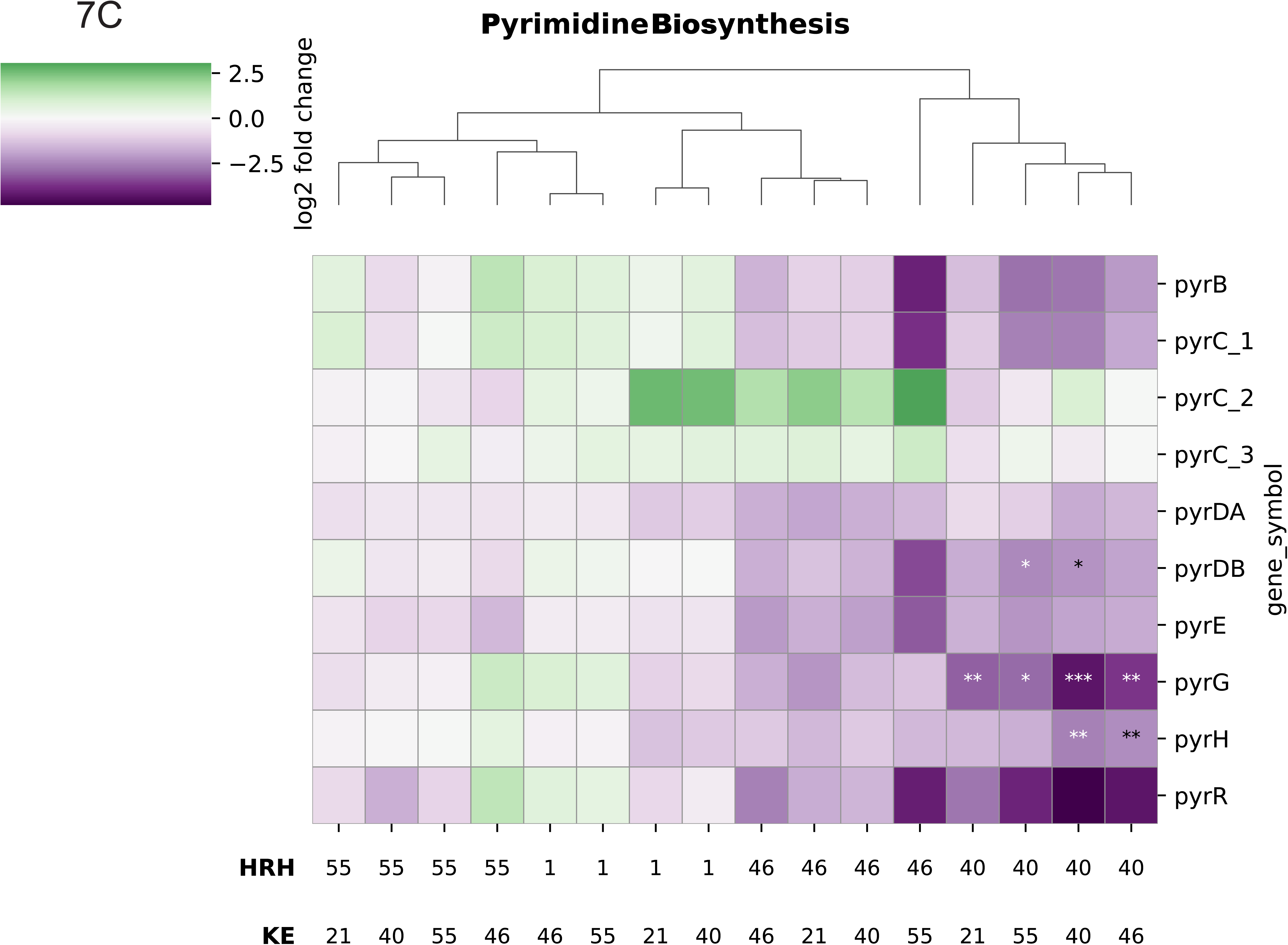
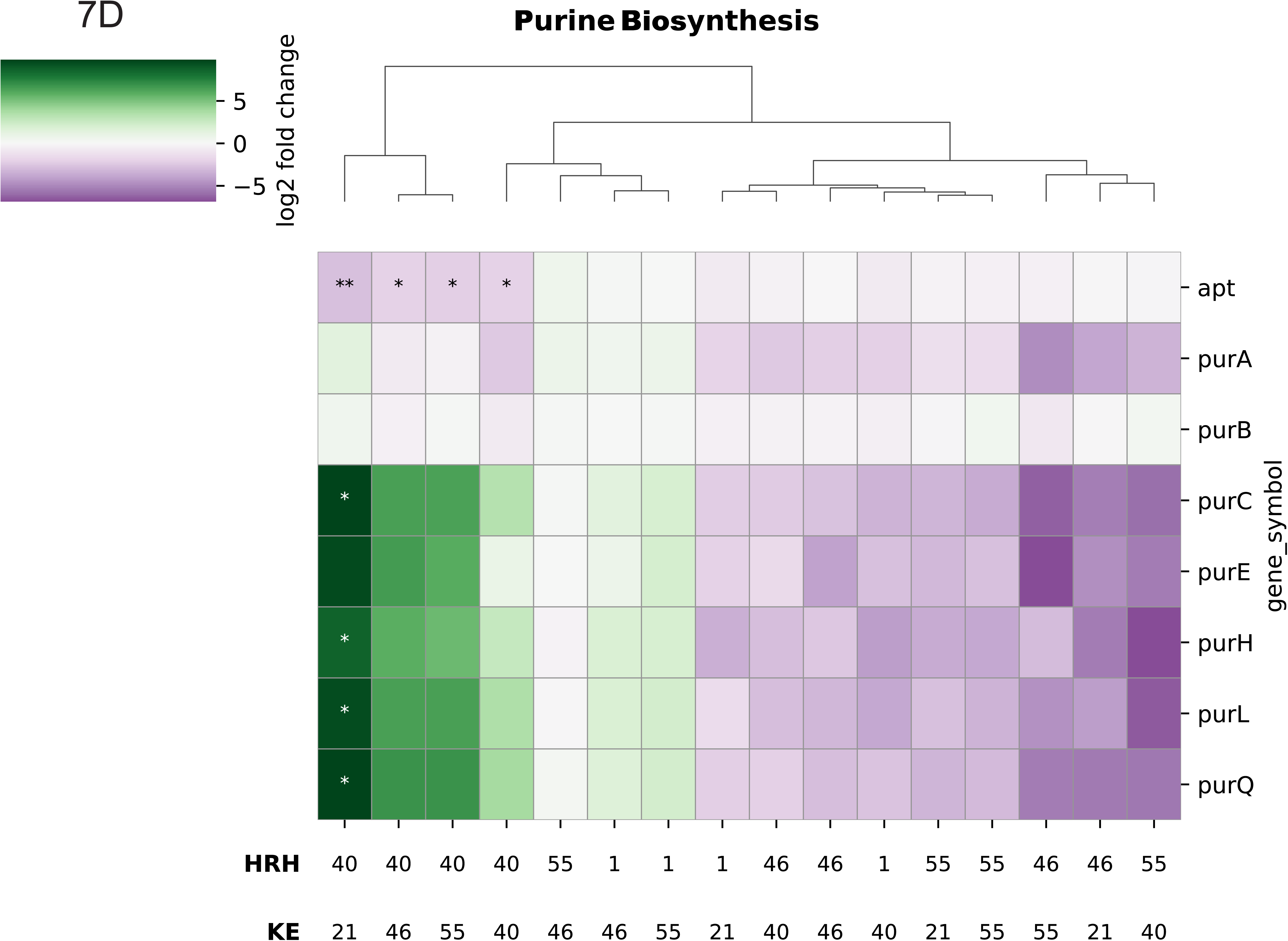
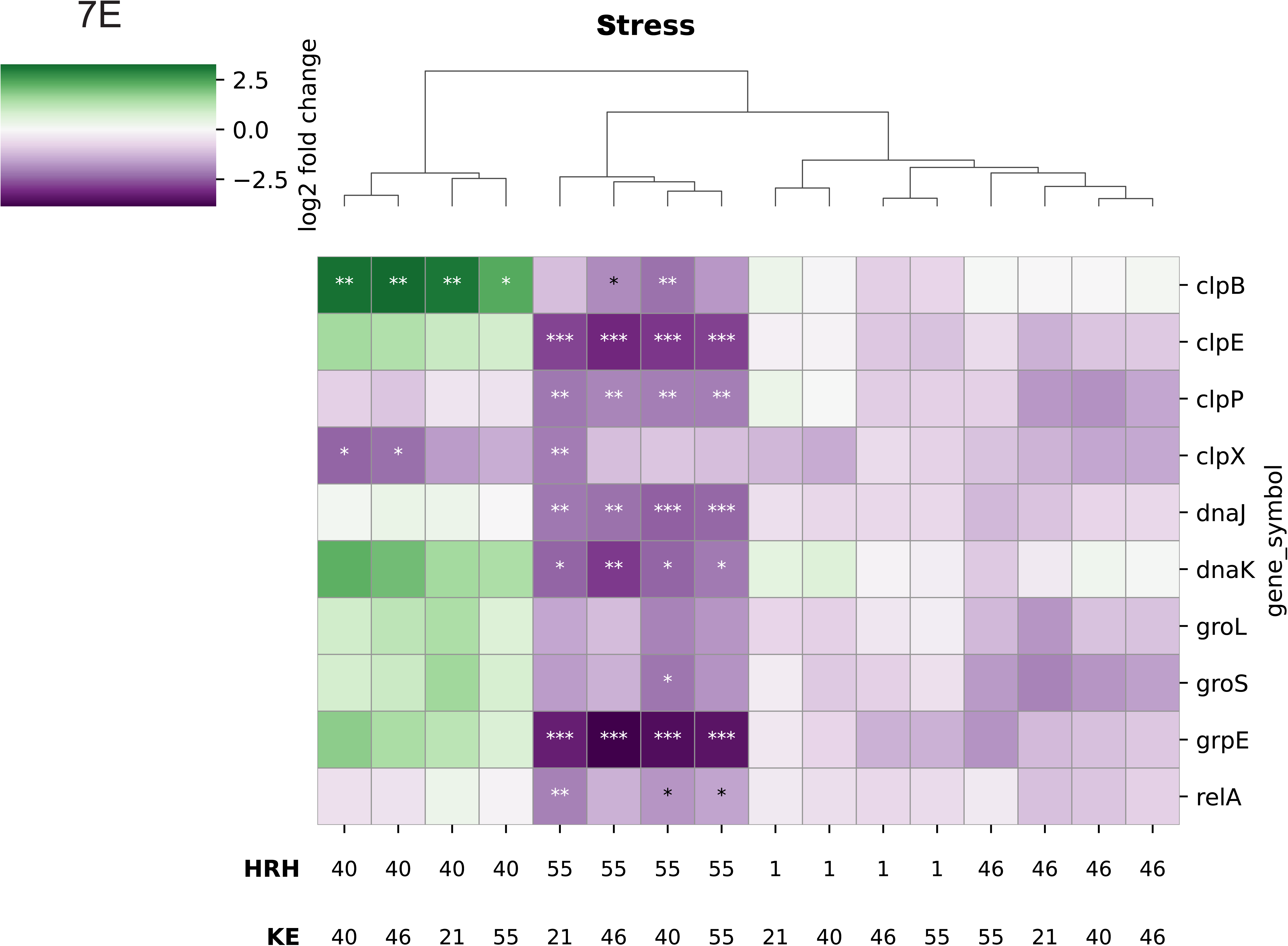
Transcript Changes in Ef. Purple depicts decreases, green depicts increases in gene expression in a co-culture compared to the respective Ef monoculture. Stars represent q-values * <0.05 **<0.01 ***<0.001. A) Ribosomal Proteins B) RNA Polymerase C) Pyrimidine Biosynthesis D) Purine Biosynthesis E) Stress

HRH46, HRH56, and OG1RF responded differently to the Ec strains. Ec reduced transcripts associated with growth rate, ribosomal proteins and RNA polymerase subunits (Fig 7A and B)), but not to the extent observed for HRH40. In contrast to HRH40, the other Ef strains had fewer transcripts for chaperone and protease genes which indicate that these Ef strains are less stressed than HRH40.

## Discussion

UTIs require high bacterial densities because of inefficient invasion into urothelial cells (26), and high densities require rapid bacterial growth in the bladder (25). Rapid growth of uropathogens was first observed in 1956 (31) and a recent assessment found a 22-minute average in vivo doubling time for uropathogens in the bladder (25). Rapid growth necessarily implies commensurate energy generation. A majority of Ec isolated from UTI patients are phylogenetic group B2 strains, although both Ec and Ef can be isolated from UTI patients (28). Recent results from urine samples from 36 UTI patients found that Ef co-isolated with eight non-B2 Ec strains, but with none of the 28 B2 Ec strains [(28) and unpublished observation]. We analyzed combinations of the three non-B2 Ec strains, their co-isolated Ef strains, the B2 strain KE21, and a standard Ef lab strain OG1RF. We utilized a nutrient-rich medium to emulate a rapid growth environment which allows both Ec and Ef to grow, although Ef grew more slowly. Our major results were that Ef had little effect on Ec growth but resulted in major transcriptome differences, the Ec-Ef interactions were not strain-specific, Ef affected the B2 strain differently, and Ef affected Ec metabolic gene transcription.

### Ef affected transcripts for genes that suggest enhanced energy metabolism

For the three non-B2 Ec strains, the four Ef strains induced Ec transcription of genes for (a) an ornithine decarboxylase, either SpeC or SpeF, which generates putrescine, (b) all enzymes of two independent pathways that degrade putrescine to succinate, and (c) all TCA cycle enzymes, except for the pyruvate dehydrogenase subunits AceE and AceF. Despite the exception, these transcript differences suggest that Ef-generated ornithine was used as an energy source.

For the B2 strain, the four Ef strains resulted in similar transcript changes with some important differences. Ef induced transcripts for genes of one or both ornithine decarboxylase and all TCA enzymes including AceE and AceF. Unlike the non-B2 strains, the B2 strain lacks the major PuuA-initiated catabolic pathway, which is a general property of B2 strains, and Ef did not induce transcripts for the minor PatA-initiated putrescine catabolic pathway, which indicates that the B2 strain does not degrade the putrescine generated from ornithine. A recently discovered putrescine homeostatic network found that intracellular putrescine positively correlates with TCA cycle enzyme transcripts (32). We propose that the putrescine content is higher in the B2 strain and that putrescine has a primarily regulatory function which includes controlling energy metabolism.

### Ef co-culture increased transcripts for glyoxylate shunt (GS) enzymes

For 13 of the 16 Ec-Ef combinations, Ef induced transcripts for GS enzymes AceA (isocitrate lyase) and AceB (malate synthase). The GS bypasses the decarboxylations that would prevent assimilation of acetate as a carbon source but is not known to be expressed during growth in either glucose-containing or nutrient-rich media. In a nutrient-limited medium, such as urine, the GS may allow greater carbon assimilation, i.e., fewer decarboxylations, but in a rich medium GS function must be different.

An alternate GS function is to reduce NADH formation which can be a problem during rapid growth if NAD+ regeneration becomes limiting. Supporting this possibility is Ef induction of transcripts in non-B2 strains for a reaction sequence that generates reduced quinones instead of NADH. The proposed reaction sequence converts malate to L-lactate to pyruvate to acetate to acetyl-CoA catalyzed by MaeB, LldD, PoxB, and AcS, respectively. Ef induced transcripts for all these enzymes at least 2-fold and sometimes much more. Both LldD, a membrane-associated FMN-dependent L-lactate dehydrogenase, and PoxB, pyruvate oxidase, generate a reduced quinone instead of NADH. A novel aspect of this sequence is MaeB-dependent non-oxidative L-lactate synthesis from L-malate instead of, or in combination with, oxidative decarboxylation. MaeB is a bifunctional oxidative decarboxylating malic enzyme/non-oxidative malolactic enzyme in *Bacillus subtilis* and Ec: the malolactic activity is favored with high NADPH (33). We propose malolactic activity because induction of LldD transcripts requires L-lactate (34). The reduced quinone-generating enzymes LldD and PoxB result in acetate formation, which can be recovered by Acs-dependent synthesis of acetyl-CoA.

For the B2 strain, Ef induced transcripts suggest a slightly different reaction sequence. Ef co-culturing increased transcripts for all TCA cycle genes, including the pyruvate dehydrogenase-encoding *aceEF*-*lpd* operon but decreased transcripts for *poxB* transcripts, which suggests preferential pyruvate conversion to acetyl-CoA instead of acetate. Because Ef induced transcripts for the energy-conserving NADH dehydrogenase and other electron transport components to a greater extent in the B2 strain, we propose that the B2 strain regenerates NAD+ better than the non-B2 strains and does not require the quinone-dependent reactions.

### Effects on macromolecular synthesis

Based on small growth rate differences, co-culturing was not expected to have large transcript changes for growth rate-associated Ec genes of macromolecular synthesis (Fig 1B). However, such changes were observed for two strain combinations. All Ef strains increased transcripts in the B2 strain KE21 for ribosomal proteins, DNA replication and topoisomerases, peptidoglycan structure, and RNA polymerase, which suggests that Ef has the potential to stimulate B2 strain growth. More than one B2 strain must be analyzed to determine if this is a general pattern. However, a recent study showed that in a biofilm incubated with an artificial urine medium, Ef stimulated growth of the B2 strain CFT073 and the Ef strain was lost after 24 hours (35). The basis for this competitive interaction could be Ef stimulation of the growth-associated genes of macromolecular synthesis, which we speculate is mediated by putrescine.

### The paradox of large transcript changes without a proportional growth rate effect

Large transcript changes were observed for metabolic and macromolecular synthesis genes without corresponding changes in growth rates. To account for this discrepancy, we note that the experiments were performed in a nutrient-rich medium which may not support a much faster growth rate. Growth rate differences may become apparent in nutrient-poor environments. The transcript changes could result from medium alterations by one of the bacterial strains, e.g., ornithine secretion by Ef. Furthermore, transcript changes do not imply protein concentration changes: for ribosomal proteins, translational as well as transcriptional control determine protein levels. Another example of a discrepancy between transcription and protein content is *glnA* which codes for glutamine synthetase. B2 strains have 60-fold higher *glnA* transcripts than non-B2 strains, but because transcription is initiated from the *glnAp1* promoter instead of the *glnAp2* promoter in B2 strains, *glnA* transcripts are not translated (36, 37). Even if transcription correlates with protein concentrations, protein activity does not necessarily change proportionally because enzyme activity is controlled by a various factors that include metabolite concentrations, allosteric control, and covalent modifications.

### What is the point of the interspecies interactions?

Our goal was to examine the interaction between co-isolated Ef and non-B2 Ec. We propose that Ef produces ornithine which is utilized as an energy source for the non-B2 EC strains. Transcript levels for metabolic enzymes are higher, which is likely a response to an energy source that is not available in our nutrient-rich GT medium, but which can be utilized simultaneously with available energy sources. The beneficial effect of Ec on Ef is not clear because of the absence of transcript changes for metabolic enzymes in Ef. One possibility is that Ec provides acetate which Ef requires (38). This possibility is consistent with Ef-induced transcription of the non-B2 Ec gene for acetate-generating pyruvate oxidase. Ef did not induce this gene in the B2 strain.

## Methods and Materials

### Culture Growth

Strains were grown overnight on species selective media, Bile Esculin Azide; which selects for *Enterococcus faecalis*, and MacConkey; which selects for *Escherichia coli*. This ensures that the cultures are mono-cultures during pre-growth. Multiple single colonies were collected and grown in 1 mL of GT media (0.5% glucose, 1% tryptone, 0.25% NaCl) for 2 hours with aeration at 250 rpm at 37°C. Strains were then diluted to 0.01 OD_600_ and mixed in 10 mL of GT media and aerated at 250 rpm for 2.5 hours (approximately 5 doublings for *E. coli*). After interacting, the cells were pelleted via centrifugation at 8000xg at 4°C, media was removed and the pellets frozen overnight at -80°C.

### RNA isolation

RNA was isolated using a protocol described previously (24). Briefly, cells were homogenized using a bead beater (MP Bio) at the speed setting (6.5) for 6 cycles of 45 seconds with 5 minute rests between each cycle on ice. RNA was isolated following the Qiagen RNeasy mini prep kit. DNA removed using DNase I (Sigma) treatment, and purified RNA concentrated using a GeneJet RNA cleanup and concentration kit (Thermo). Only RNA with a Qbit RNA IQ score of 8 or greater and OD 260/280 values greater than 2.0 were selected for RNAseq.

### RNA sequencing

Purified RNA was submitted to the University of Texas at Dallas Genome Center for sequencing using an Illumina Nextseq 2000 single end 150 bp read length to a target depth of 90 million reads per sample.

### RNA analysis

Prokka-annotated genomes (39) were clustered using CD-HIT (40) then used to establish a metatranscriptome by merging all of the gene calls from the annotations at 95% sequence identity. Kallisto (41) was then used to align/quantify reads against the metatranscriptome, and sleuth (42) to perform differential expression calling. Separate sleuth analysis for each strain of interest was performed, along with differential expression calling for each co-culture of that strain with a representative of the other species (e.g., focusing on a single *E. coli* strain, its interaction with each *E. facaelis* strain) relative to the corresponding monoculture.

### Growth for OD Measures

Cells were pre-grown as described above. Cells were then diluted and then 150 µL of culture were transferred to a 96-well plate, and grown in a Epoch plate reader (BioTek) aerated at the highest shaking setting at 37°C for 2.5 hours with optical measures taken every 30 minutes at OD_600_.

### Strains

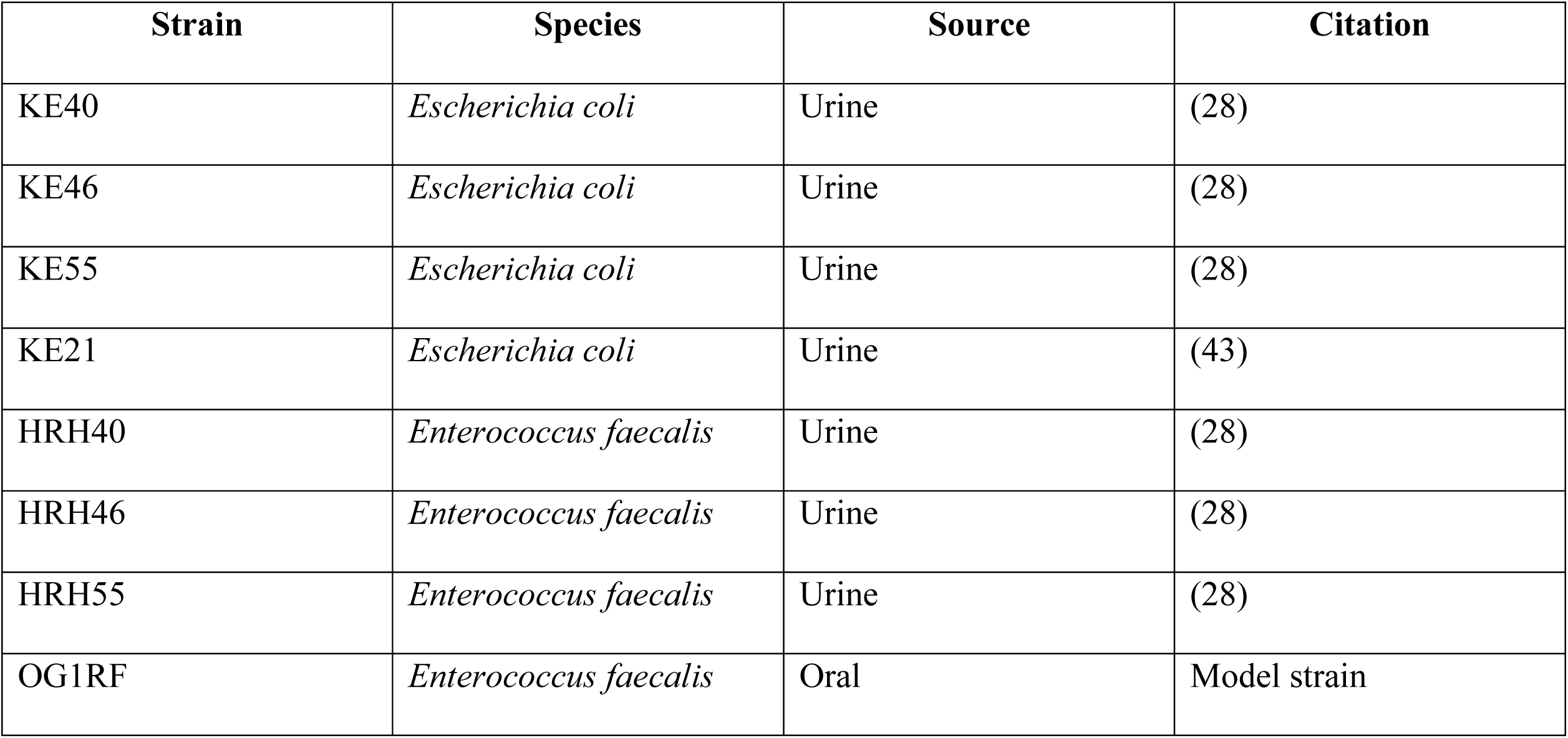

| Strain | Species | Source | Citation |
| --- | --- | --- | --- |
| KE40 | *Escherichia coli* | Urine | (28) |
| KE46 | *Escherichia coli* | Urine | (28) |
| KE55 | *Escherichia coli* | Urine | (28) |
| KE21 | *Escherichia coli* | Urine | (43) |
| HRH40 | *Enterococcus faecalis* | Urine | (28) |
| HRH46 | *Enterococcus faecalis* | Urine | (28) |
| HRH55 | *Enterococcus faecalis* | Urine | (28) |
| OG1RF | *Enterococcus faecalis* | Oral | Model strain |

## Acknowledgments

This work was supported by NIH R35 GM128637 (to L.F.).

